# Metagenomic analysis reveals the abundance and diversity of opportunistic fungal pathogens in the nasopharyngeal tract of COVID-19 patients

**DOI:** 10.1101/2022.02.17.480819

**Authors:** M. Nazmul Hoque, M. Shaminur Rahman, Md. Murshed Hasan Sarkar, Md Ahashan Habib, M. Anwar Hossain, M. Salim Khan, Tofazzal Islam

## Abstract

The nasopharyngeal tract (NT) of human is a habitat of a diverse microbial community that work together with other gut microbes to maintain the host immunity. In our previous study, we reported that SARS-CoV-2 infection reduces human nasopharyngeal commensal microbiome (bacteria, archaea and commensal respiratory viruses) but increases the abundance of pathobionts. This study aimed to assess the possible changes in the resident fungal diversity by the inclusion of opportunistic fungi due to the infection of SARS-CoV-2 in the NT of humans. Twenty-two (n = 22) nasopharyngeal swab samples (including COVID-19 = 8, Recovered = 7, and Healthy = 7) were collected for RNAseq-based metagenomics analyses. Our results indicate that SARS-CoV-2 infection significantly increased (p < 0.05, Wilcoxon test) the population and diversity of NT fungi with a high inclusion of opportunistic pathogens. We detected 863 fungal species including 533, 445, and 188 species in COVID-19, Recovered, and Healthy individuals, respectively that indicate a distinct microbiome dysbiosis due to the SARS-CoV-2 infection. Remarkably, 37% of the fungal species were exclusively associated with SARS-CoV-2 infection, where *S. cerevisiae* (88.62%) and *Phaffia rhodozyma* (10.30%) were two top abundant species in the NT of COVID-19 patients. Importantly, 16% commensal fungal species found in the Healthy control were not detected in either COVID-19 patients or when they were recovered from the COVID-19. Pairwise Spearman’s correlation test showed that several altered metabolic pathways had significant positive correlations (r > 0.5, p < 0.01) with dominant fungal species detected in three metagenomes. Taken together, our results indicate that SARS-CoV-2 infection causes significant dysbiosis of fungal microbiome and alters some metabolic pathways and expression of genes in the NT of human. Findings of our study might be helpful for developing microbiome-based diagnostics, and also devising appropriate therapeutic regimens including antifungal drugs for prevention and control of concurrent fungal coinfections in COVID-19 patients.

**Author summary:** The SARS-CoV-2 is a highly transmissible and pathogenic betacoronavirus that primarily enters into the human body through NT to cause fearsome COVID-19 disease. Recent high throughput sequencing and downstream bioinformatic analyses revealed that microbiome dysbiosis associated with SARS-CoV-2 infection are not limited to bacteria, and fungi are also implicated in COVID-19 development in susceptible individuals. This study demonstrates that SARS-CoV-2 infection results in remarkable depletion of NT commensal fungal microbiomes with inclusion of various opportunistic fungal pathogens. We discussed the role of these altered fungal microbiomes in the pathophysiology of the SARS-CoV-2 infection. Our results suggest that dysbiosis in fungal microbiomes and associated altered metabolic functional pathways (or genes) possibly play a determining role in the progression of SARS-CoV-2 pathogenesis. Thus, the identifiable changes in the diversity and composition of the NT fungal population and their related genomic features demonstrated in this study might lay a foundation for better understanding of the underlying mechanism of co-pathogenesis, and the ongoing development of therapeutic agents including antifungal drugs for the resolution of COVID-19 pandemic.

## Introduction

Coronavirus disease (COVID-19), emerged as one of the deadliest human diseases, is considered as the fast expanding pandemics since the 1918 Spanish flu with serious consequences for global health and economy [1–3]. Since SARS-CoV-2 emerged in the human population, the global scientific community is working round the clock to find good strategies for the containment and treatment of this pandemic virus, SARS-CoV-2 [4]. Upon inhalation, SARS-CoV-2 primarily enters the nasal epithelial cells of the human NT through the ACE2 and TMPRSS2 receptors [5], and then gradually move towards the lung to initiate infection followed by onset of acute respiratory distress syndrome (ARDS) [6]. Viral replication in the nasopharyngeal epithelial cells elicits direct adverse effects on resilient microbiomes [4,7,8], and induces local immune cells to quickly and abundantly secrete cytokines and chemokines [9]. Subsequently, severe lung damage and immunologic derangement resulting from SARS-CoV-2 infection or its treatment predispose to coinfections with multiple pathogens, including bacteria, other viruses and fungi [7,10,11]. Despite the potentially poor prognosis, data on SARS-CoV-2 coinfection is still scarce and thus, the universal pervasiveness of coinfection among COVID-19 patients is unknown [7, 10]. Clinical trials and high throughput sequencing (metagenomic and RNAseq)-based investigations on SARS-CoV-2 revealed that severely and non-severely ill COVID-19 patients had coinfections with respiratory viral pathogens [12], and bacteria and/or fungi [4,7,10,13]. A retrospective study found that the coinfection rate of SARS-CoV-2 and influenza virus was as high as 57.3% in COVID-19 patients during the outbreak period in Wuhan [14]. The coinfection in COVID-19 patients may be a predisposing factor of increased morbidity and mortality rates throughout the globe [10,12,15]. It is well known that viral respiratory infection such as influenza can be complicated by bacterial and fungal coinfections [16]. Previously, the SARS outbreak was characterised by an high rate of nosocomial transmission of drug-resistant microorganisms [7, 16].

Fungal infections are known to be among the infectious complications related to the damage caused by viral pulmonary infections, particularly in patients admitted to intensive care units with ARDS [10]. Patients with severe COVID-19 have also emerged as a population with a high risk of fungal infections [10, 17]. Opportunistic fungal infections following severe respiratory viral illness has been described most frequently with influenza complicated by respiratory failure, with an incidence of invasive pulmonary aspergillosis [18, 19]. There are reports that immunocompromised COVID-19 patients were at a higher risk of development of mysterious fungal infection known as mucormycosis or black fungus [20, 21]. Recently other non-Aspergillus fungal coinfections, including mucormycosis in India, have been reported in those with severe COVID-19 pulmonary disease [22]. Meanwhile, a descriptive study held by Chen et al. (2020) showed that the coinfected fungi includes *Aspergillus* spp., *Candida albicans*, and *Candida glabrata* [23]. Fungal coinfection was the main cause of death for SARS patients, accounting for 25–73.7% in all causes of death [24]. Besides, in the past decade, increasing reports of severe influenza pneumonia resulting in ARDS complicated by fungal infection were published [25]. With the aggravating pathogenesis of SARS-CoV-2, most of the COVID-19 patients usually undergone to the in-time use of broad spectrum antibiotics, dexamethasone, and immunosuppressive therapies with corticosteroids or immunomodulators for bacterial coinfections [17, 26], while the diagnosis of fungal coinfection is always delayed or neglected. Based on the experience of SARS in 2003 and the cases of invasive aspergillosis combined with severe influenza, it is critically important to pay attention to the probability of COVID-19 accompanied by fungal infections. However, as for fungal coinfection in COVID-19 patients, only few studies have reported it, which may have been neglected. Clinically, many COVID-19 patients did not undergo fungal assessment at the beginning, moreover, it is difficult to detect fungus with a single sputum fungal culture [24].

Human microbiota plays a critical role in immunity and health of individual hosts, and thus, microbiome dysbiosis in the respiratory tract by the pathogenic respiratory virus like SARS-CoV-2 can increase the mortality rate in patients [7,27,28]. Coinfection can also change the upper airway microbiome homeostasis and thus, triggers the infection and stimulates immune cells to produce more severe inflammation [7,10,29]. Therefore, we hypothesize that during this migration, propagation and immune response, the inhabitant fungal microbiomes in the respiratory airways are altered or dysbiosed, and inclusion of some of the fungal pathobionts might aggravate the progression and lethality in COVID-19 patients. More importantly, identification of the coinfecting fungal pathobiomes may also lead to a new strategy for treatment of COVID-19 patients. Therefore, timely diagnosis of coinfecting fungal pathogen(s) in COVID-19 patients is important to initiate appropriate therapy and limit the overuse of antimicrobial agents. To shed light on the effects and consequences of SARS-CoV-2 infections on changes in the resident fungal microbiota in the NT, we conducted a high throughput RNAseq analysis of the nasopharyngeal swabs randomly collected from Healthy controls, COVID-19 patients and COVID-19 Recovered inviduals. Our results demonstrate that SARS-CoV-2 infection is critical for inclusion of opportunistic fungal microbiomes with loss of salutary fungi in the NT. Besides, we conducted a comparative metabolic functional analysis to identify the potential biological mechanisms linking the shift of fungal population in NT, and their correlation with differentially abundant fungal taxa in the COVID-19 patients, Recovered humans and Healthy controls. Taken together, our data suggest a strong and critical association of altered fungal microbiome in the pathophysiology of SARS-CoV-2 infections in the nasal cavity of COVID-19 patients.

## Results

In order to detect the dysbiosis of the inhabitant fungal microbiomes in the NT after SARS-CoV-2 infection, we analyzed 22 nasal swab samples from Healthy individuals, COVID-19 patients and Recovered humans. The demographics, health-related characteristics, and symptomatology of the study subjects are described in S1Table. Overall, 15 study people were male (68.18%) and seven were female (31.82%) with a median age of 40.87 years. In this study, patients were diagnosed positive for SARS-CoV-2 infections (COVID-19) on an average 5.6 days after the onset of pneumonia like clinical symptoms, and these patients tested negative for SARS-CoV-2 (Recovered) on an average 15.12 days after the initial COVID-19 confirmatory diagnosis. The confirmed COVID-19 patients received medication for an average period of 15.71 days. The Healthy control subjects however did not show any signs and symptoms of respiratory illness (S1 Table). We were able to characterize both common and differentially abundant fungal taxa in each sample groups along with concurrent metabolic functional perturbations.

### Fungal diversity in the nasal cavity is affected by SARS-CoV-2 infection

To understand whether SARS-CoV-2 infection alters the composition and diversity of the NT fungal microbiomes, we examined both within sample (alpha) and across the samples (beta) diversities of the detected fungal community in Healthy human, COVID-19 patients and Recovered humans (Fig 1). The alpha diversity measured using Observed, Chao1, ACE, Shannon, Simpson and InvSimpson indices showed significant differences in fungal community richness, keeping substantially higher diversity in the microbial niche of COVID-19 (p = 0.01; Wilcoxon test) followed by Recovered (p = 0.05; Wilcoxon test) and Healthy (p > 0.05; Wilcoxon test) samples (Fig 1). The Bray–Curtis dissimilarity distance estimated principal coordinate analysis (PCoA) plot showed that fungal composition in the NT nasal cavity differed according to sample categories of the study people (Fig 2). The beta diversity of microbiomes however did not vary significantly (p > 0.05, PERMANOVA test) according to the sex (male or female) of the study people (Fig 2).

**Fig 1.**
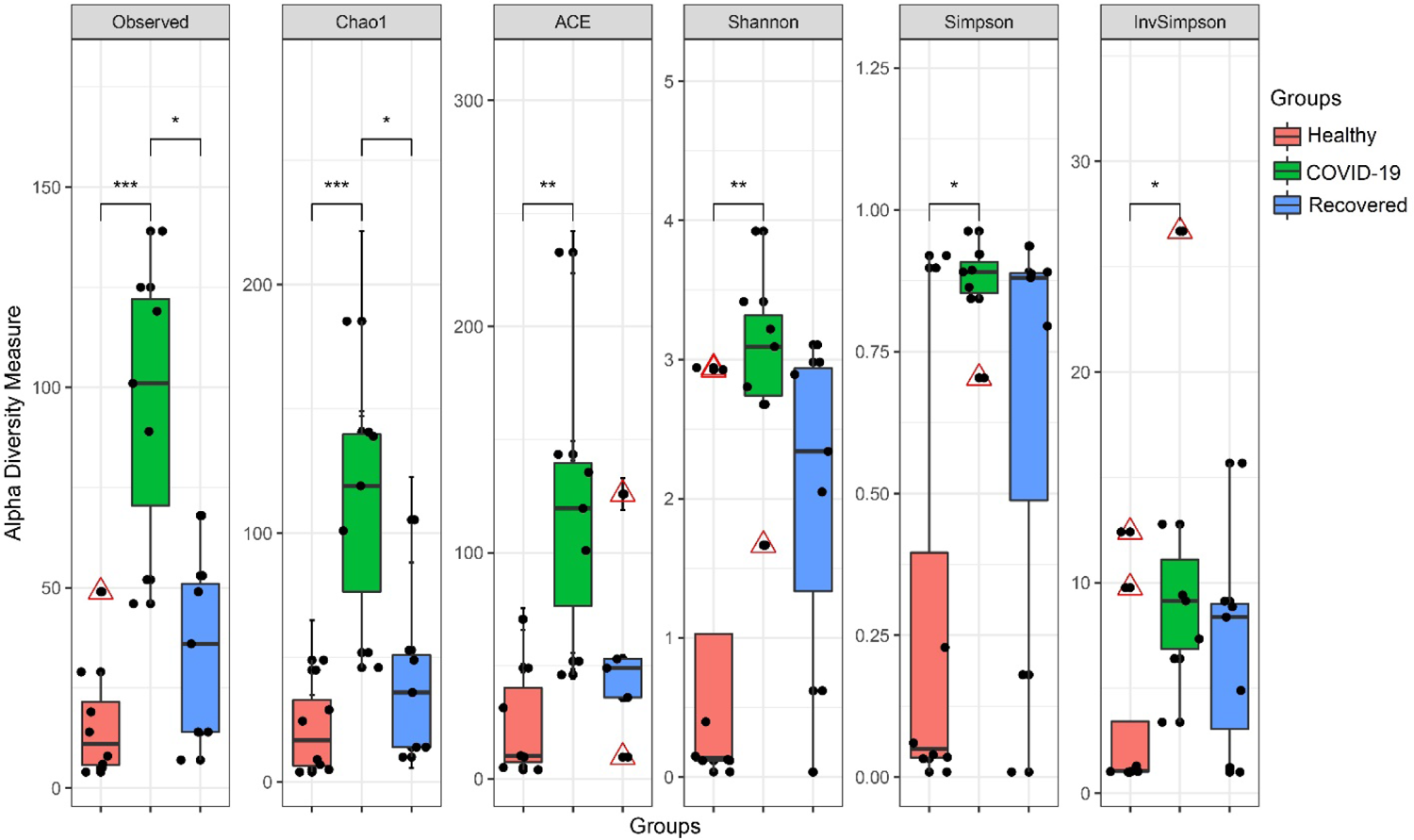
Within subject (Alpha) diversity measure. Observed species, Chao1, ACE, Shannon, Simpson and InvSimpson indices estimated within sample fungal diversity of Healthy, COVID-19 and Recovered cases are plotted on boxplots and comparisons are made with pairwise Wilcoxon rank sum tests. Significance level (p-value) 0.0001, 0.001, 0.01, and 0.05 are represented by the symbols “****”, “***”, “**”, and “*”, respectively.

**Fig 2.**
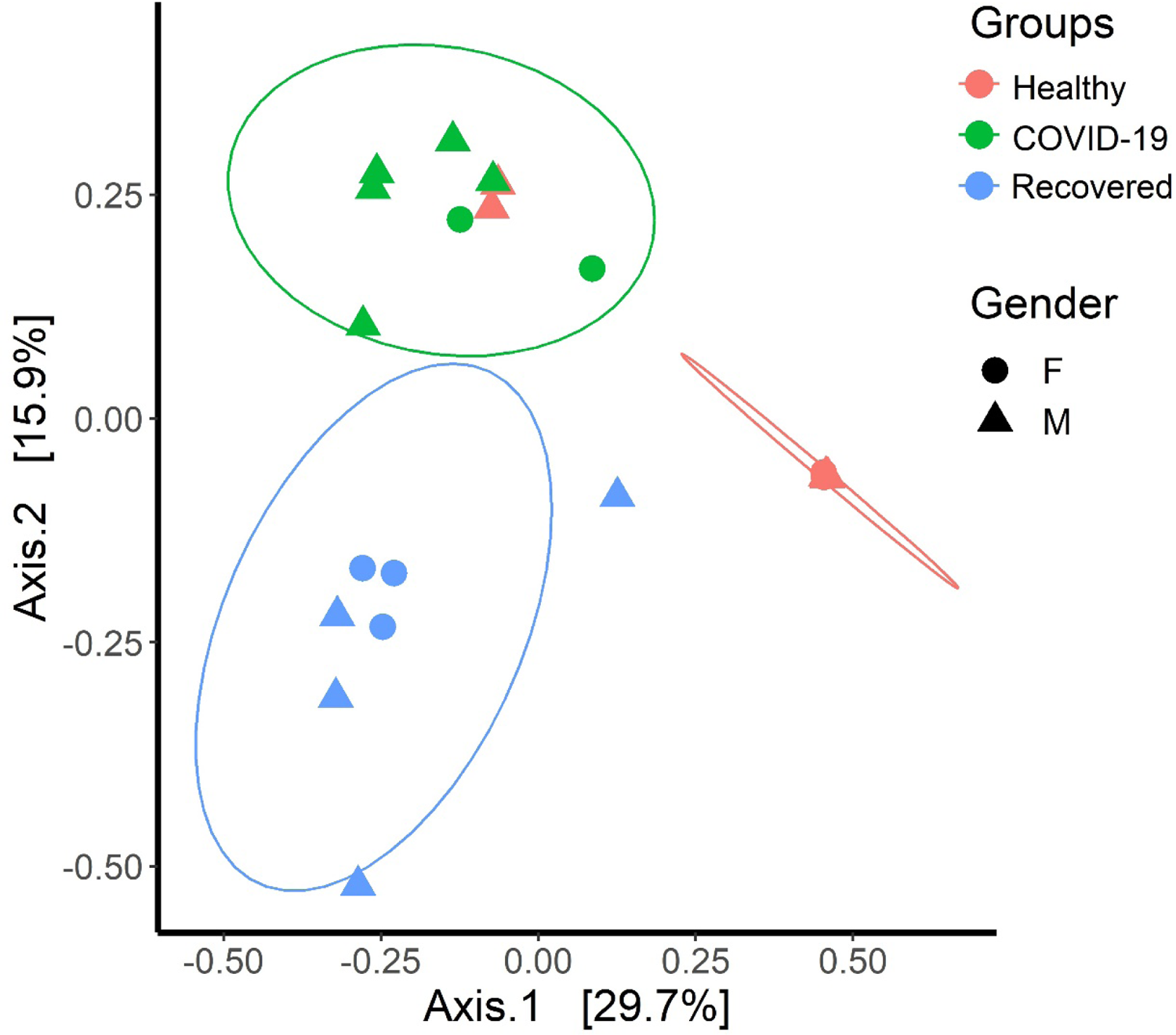
Between subject (Beta) diversity measure. Fungal beta diversity is measured with Bray-Curtis dissimilarity distances and visualized on principal coordinate analysis (PCoA) plot. The samples are colored according to subject groups (e.g., red: Healthy, green: COVID-19 and blue: Recovered) and joined with the respective ellipses. The shapes represent the gender of the assigned subjects: circular for female (F) and triangular for male (M). Pairwise comparisons on a distance matrix using PERMANOVA test under reduced model shows significant fungal community differences among the groups (p < 0.01, PERMANOVA test).

The unique and shared distribution of fungal taxa found in the three metagenome groups is presented by comprehensive Venn diagrams (Fig 3). A total of 28 fungal phyla were detected in three metagenomes including 16, 24 and 18 phyla in Healthy, COVID-19 and Recovered samples, respectively. Among these phyla, six phyla had sole association with SARS-CoV-2 infections, and 10 were found to be shared across three sample groups (Fig 3A, S1 Data). In this study, we detected 190 orders of fungi including 78, 131 and 104 in Healthy, COVID-19, and Recovered nasopharyngeal samples, respectively, and of them, 52 orders had sole association with SARS-CoV-2 infection, and only 35 were common in three sample groups (Fig 3B, S1 Data). Likewise, 532 fungal genera were identified, of which 57, 213 and 128 genera had sole association with Healthy, COVID-19, and Recovered subjects, respectively, and only 34 genera were found to be shared across three metagenomes (Fig 3C, S1 Data). One of the noteworthy findings of the present study was the detection of 862 fungal species and of them, 188, 533 and 445 species were found in Healthy, COVID-19, and Recovered samples, respectively. However, among the detected fungal species, only 65 (7.54%) were found to be shared across the given conditions (Fig 3D). Remarkably, compared to Healthy controls and Recovered cases, COVID-19 patients had sole association of higher number of fungal species (n=315, 36.54%) which probably due to the opportunistic inclusion during the pathogenesis of SARS-CoV-2 infection. Similarly, Recovered humans swab samples had sole association of 227 (26.33%) fungal species revealing the re-establishment beneficial commensal flora after the recovery of SARS-CoV-2 infections. Conversely, 81 fungal species had sole association with healthy states (Healthy control) of the humans (Fig 3D), and none of these species was detected in the COVID-19 patients swab samples demonstrating that these commensal microbes underwent to dysbiosis through the effect of SARS-CoV-2 infection (Fig 3D, S1 Data).

**Fig 3.**
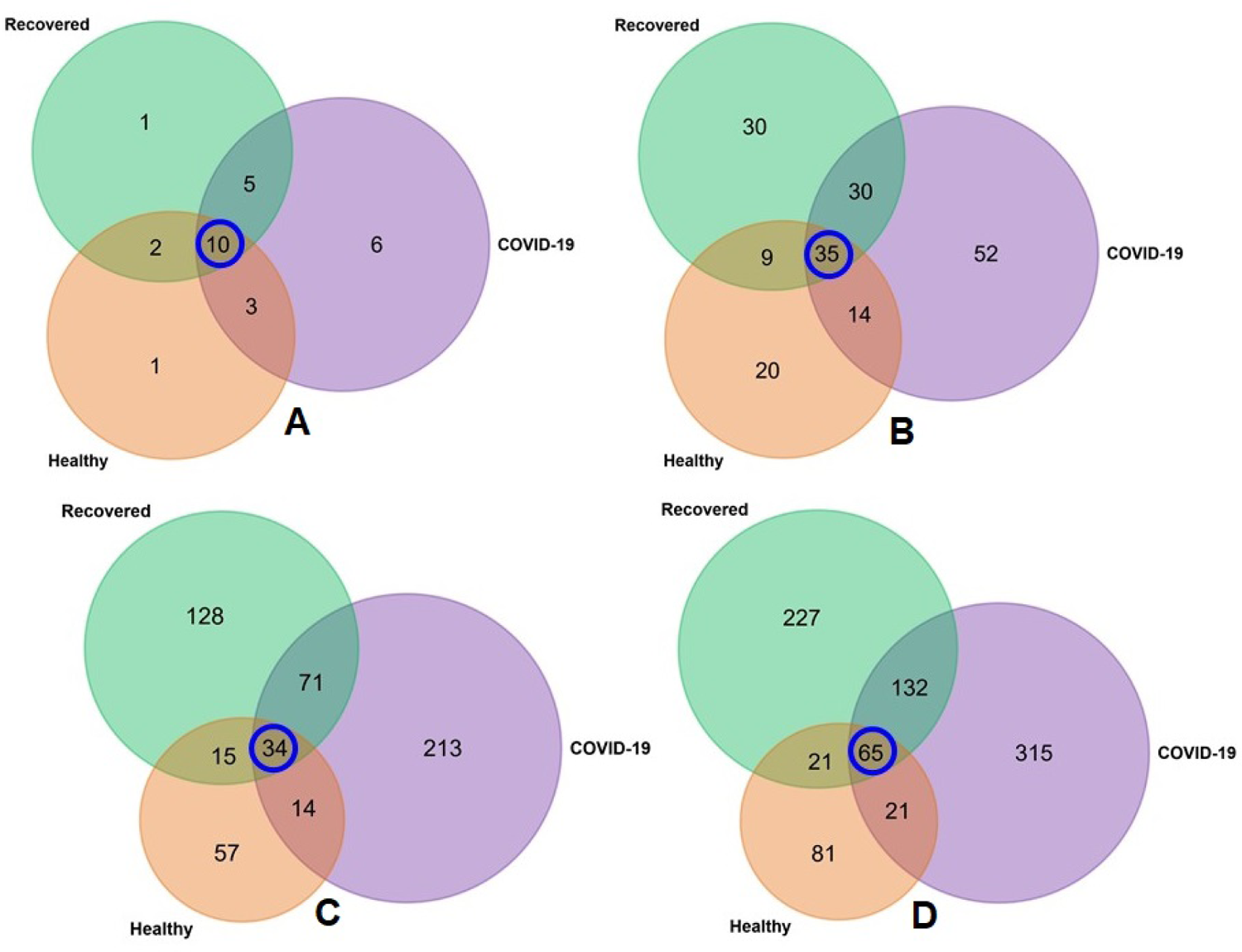
Taxonomic composition of fungal microbiomes. Venn diagrams representing the unique and shared fungal taxa in Healthy, COVID-19, and Recovered nasopharyngeal sample groups. (A) Venn diagram showing unique and shared fungal phyla. Out of 28 detected phyla, only 10 phyla (highlighted in blue circle) were found to be shared in the metagenomes. (B) Venn diagram comparison of 190 orders of fungi detected across the sample groups, of which only 35 (highlighted in blue circle) orders were found to be shared among the conditions. (C) Venn diagrams representing unique and shared fungal genera identified in three metagenomes. Of the detected fungal genera (n=532), 57, 213 and 128 genera had sole association with Healthy, COVID-19, and Recovered subjects, respectively, and only 34 genera (highlighted in blue circle) were found to be shared across three metagenomes. (D) Venn diagrams representing unique and shared fungal species identified in three metagenomes. Of the detected fungal species (n=862), the Healthy, COVID-19 and Recovered cases had sole association of 81, 227 and 315 species, respectively, and 65 species (highlighted in blue circle) were found to be shared across the study sample groups. More information on the taxonomic result is also available in S1 Data.

### SARS-CoV-2 infection induces nasopharyngeal fungal microbiome dysbiosis

To determine whether SARS-CoV-2 infection dysbiosis the NT fungal microbiomes, we characterized fungal taxa at different taxonomic ranks (phylum to species level) across three metagenomes. In this study, fungal microbiota in COVID-19 patients, Recovered humans and Healthy control samples were predominated by *Ascomycota* (> 87.0%) phylum, however, other abundant phyla were *Basidiomycota* (2.26 to 3.83%), *Streptophyta* (1.41 to 2.20%), and *Mucoromycota* (2.06%) (Fig 4). Rest of the fungal phyla detected in all of the metagenomes had relatively lower abundances (< 1.0%) (S1 Data). Moreover, the average phyla distribution in SARS-CoV-2 infection associated metagenomes (COVID-19 and Recovered) was different compared to that in Healthy controls. Notably, the distribution phyla in COVID-19 and Recovered cases demonstrated greater similarities than those detected in Healthy controls (Fig 4, S1 Data).

**Fig 4.**
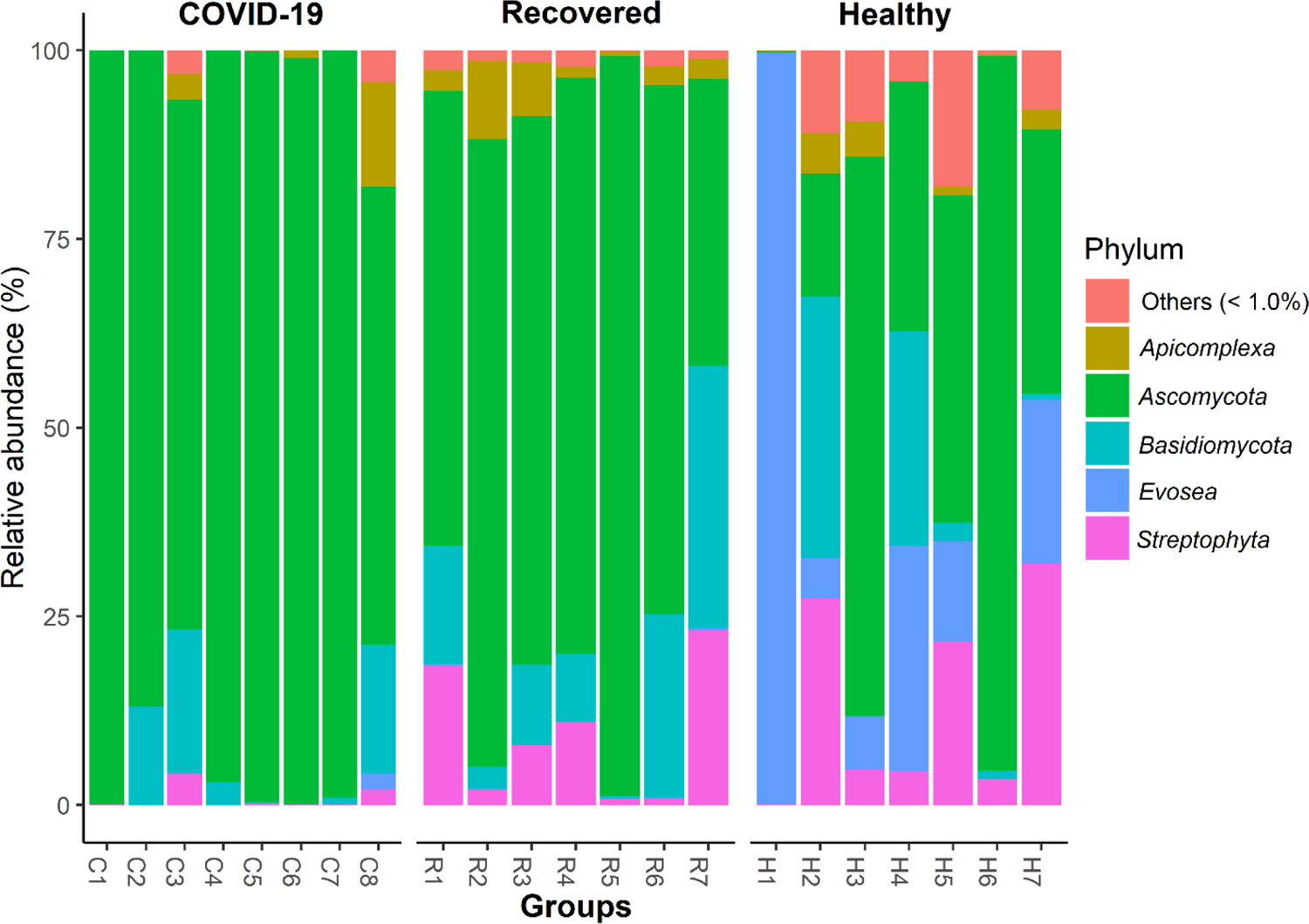
Major fungal phyla detected. The phylum-level taxonomic profile of fungal microbiomes in COVID-19 (C1-C8), Recovered (R1-R7) and Healthy (H1-H7) nasopharyngeal samples. Phyla with > 1% mean relative abundance are represented by different color codes against respective sample groups. Others (< 1%) indicates the rare taxa in each group, with mean relative abundance of < 1%.

We then pairwise compared the NT microbial consortia between the subjects with SARS-CoV-2 infection (COVID-19) and without SARS-CoV-2 infection (Healthy and Recovered) on a distance matrix using PERMANOVA test under reduced model which showed significant differences (p < 0.01, PERMANOVA test) in fungal community across the study groups. Pairwise Wilcoxon tests identified that five phyla (*Ascomycota*, *Streptophyta*, *Tubulinea*, *Bacillariophyta* and *Evosea*) were significantly different (p < 0.05, Wilcoxon test) in the Healthy, COVID-19 and Recovered metagenomes (Fig 5). Healthy and Recovered samples had significantly higher (p = 0.01, Wilcoxon test) relative abundance of *Streptophyta* than COVID-19 samples (p = 0.05, Wilcoxon test). In addition to *Ascomycota* (96.70%), the COVID-19 samples had significantly higher (p < 0.05, Wilcoxon test) relative abundance of *Tubulinea* and *Bacillariophyta* (Fig 5). The Recovered metagenome had significantly higher relative abundance of *Mucoromycota* (2.06%) and *Apicomplexa* (0.32%) compared to Healthy controls and COVID-19 patients (≤ 0.01% in both). Conversely, the Healthy humans NT swab samples had higher relative abundances of *Cnidaria*, *Chlorophyta*, *Discosea*, *Evosea* and *Rhodophyta* compared to COVID-19 and Recovered samples (Fig 5, S1 Data).

**Fig 5.**
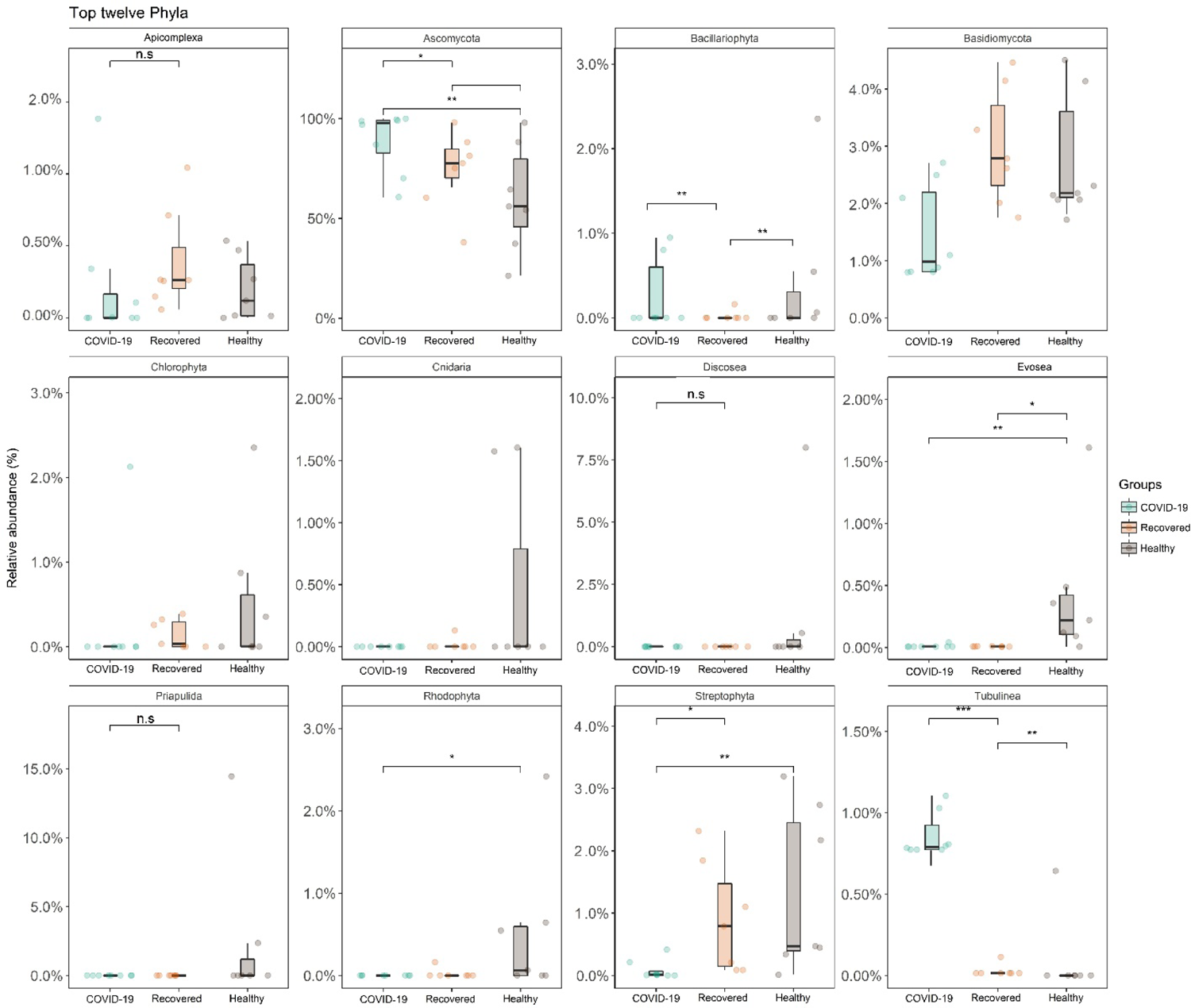
Top twelve fungal phyla detected. The phylum-level taxonomic abundance of fungal microbiomes in COVID-19, Recovered and Healthy nasopharyngeal samples. The diversity for each phylum is plotted on boxplots and comparisons are made with pairwise Wilcoxon test rank sum tests. Significance level (p-value) 0.0001, 0.001, 0.01, 0.05, and 0.1 are represented by the symbols “****”, “***”, “**”, “*”, and “n.s”, respectively.

The present microbiome study demonstrated notable differences in both composition and the relative abundances of fungal taxa at genus-level among COVID-19 patients, Recovered humans and Healthy controls. The relative abundances of the top 15 fungal genera were compared among the Healthy, COVID-19 and Recovered cohorts (Fig 6). Among these predominating genera, *Nannochloropsis* (81.58%) was the top abundant genus in Healthy controls while *Saccharomyces* (96.49%) and *Aspergillus* (81.90%) were the predominating genera in COVID-19 patients and Recovered humans NT swabs, respectively (Fig 6, S1 Data).

**Fig 6.**
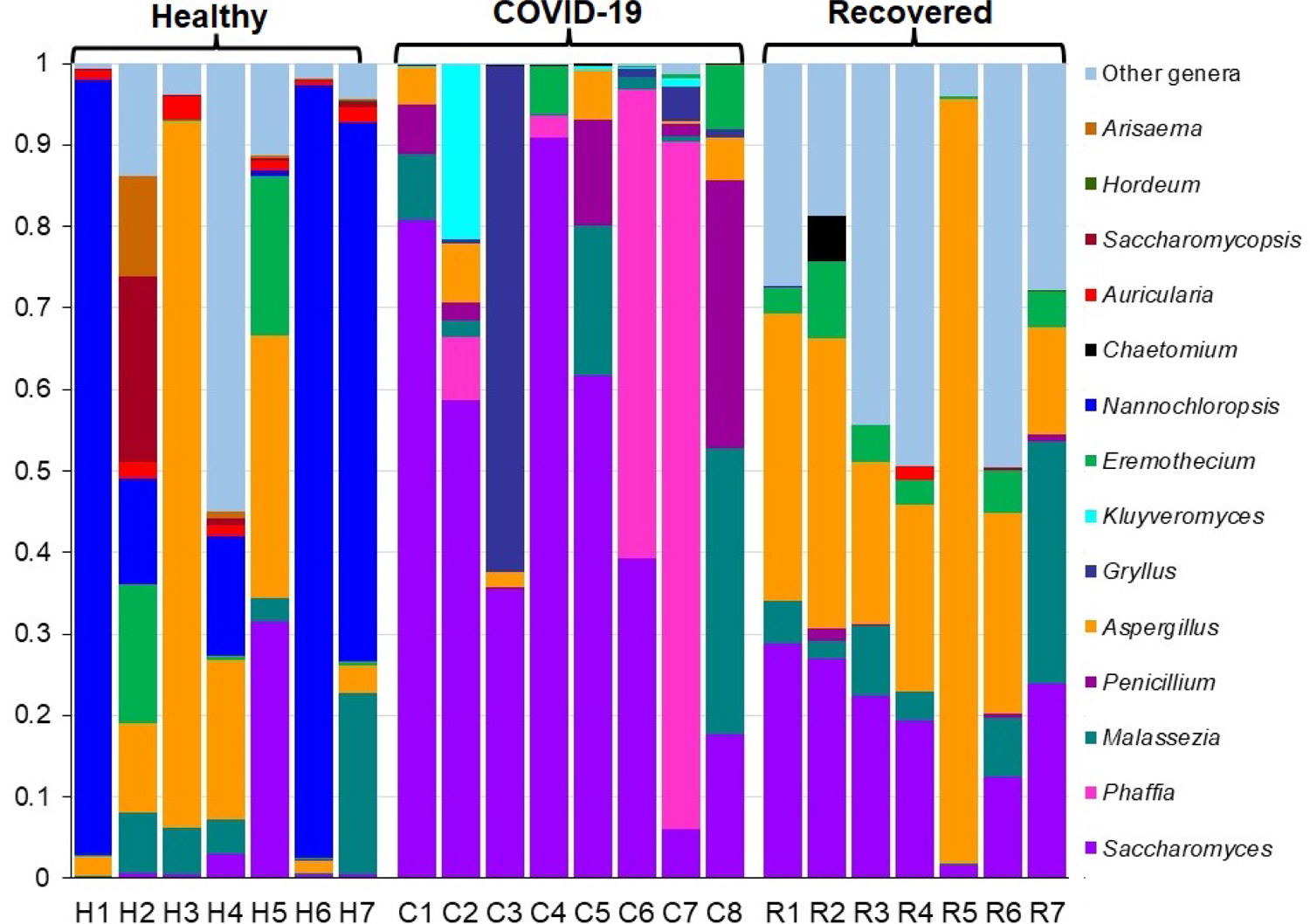
The genus-level taxonomic profile of fungal microbiomes. The bar plots representing the relative abundance of 15 top abundant fungal genera in Healthy (H1-H7), COVID-19 (C1-C8), and Recovered (R1-R7) human nasopharyngeal samples. Fourteen genera are sorted from bottom to top by their deceasing proportion of the mean relative abundances, with the remaining genera keeping as ‘Other genera’. Each stacked bar plot represents the abundance of fungal genera in each sample of the corresponding category. Notable differences in fungal populations are those where the taxon is abundant in COVID-19 and Recovered samples, and effectively not detected in the Healthy controls. The distribution and relative abundance of the fungal genera in the study metagenomes are also available in S1 Data.

The other predominant fungal genera in Healthy controls were *Aspergillus* (3.19%), *Saccharomyces* (2.60%), *Triticum* (2.17%), *Auricularia* (1.30%) and *Eremothecium* (1.10%). Conversely, *Phaffia* (2.88%) in COVID-19 patients, and *Saccharomyces* (4.71%), *Malassezia* (3.49%), *Triticum* (1.69%), and *Eremothecium* (1.20%) in Recovered humans were other abundant fungal genera. Though rest of the genera had relatively lower abundances (<1.0%), but their relative abundances differed across three sample groups (Fig 6, S1 Data). Fungal genera identified in the Healthy humans NT swabs resemble more similarity to those detected in Recovered humans NT swab when compared with those detected in COVID-19 patients NT swabs (S1 Data).

### Differentially abundant and altered fungal species are correlated with SARS-CoV-2 pathophysiology

To examine whether species level composition and relative abundance of the fungal taxa statistically differ across the sample groups, we examined pairwise Spearman correlation of abundance of all taxa identified. This differential analysis revealed that genus level microbiome composition and diversity discrepancy was more evident at species level. However, presence of few predominating fungal species in each sample category suggested that the crucial differences might be found at the strain level. The Healthy controls nasal swab samples were dominated by *Nannochloropsis oceanica* (47.93%), *Saccharomyces pastorianus* (34.42%), *Saccharomyces cerevisiae* (2.80%), *Aspergillus pseudoglaucus* (1.84%), *Aspergillus penicillioides* (1.25%), *Paecilomyces variotii* (1.24%), and *Eremothecium gossypii* (1.06%) (Fig 7, S1 Data). Despite, 36.54% of the fungal species were exclusively associated with SARS-CoV-2 infection, only *S. cerevisiae* (88.62%) was detected as the most predominating species in COVID-19 patients. However, other abundant species identified in this metagenome included *Phaffia rhodozyma* (10.30%), *S. pastorianus* (0.43%), *Paecilomyces variotii* (0.37%), and *A. pseudoglaucus* (0.17%) (Fig 7, S1 Data). Rest of the species had relatively lower abundances (< 0.1%) in COVID-19 metagenome and possibly played an opportunistic role in the SARS-CoV-2 pathogenesis (S1 Data). In addition, Recovered humans NT swab samples were mostly dominated by different species of *Aspergillus* genus (> 80.0%) such as *A. penicillioides* (36.64%), *A. keveii* (23.36%), *A. oryzae* (10.05%), *A. pseudoglaucus* (4.42%), *A. flavus* (1.44%), *A. fumigatus* (1.34%), *A. glaucus* (1.16%) and *A. lentulus* (1.10%). However, other dominating fungal species in this metagenome were *S. cerevisiae* (4.66%), *Malassezia restricta* (2.63%), *E. gossypii* (1.20%), *P. variotii* (1.08%), and rest of the genera had relatively lower abundances (< 1.0%) (Fig 7, S1 Data).

**Fig 7.**
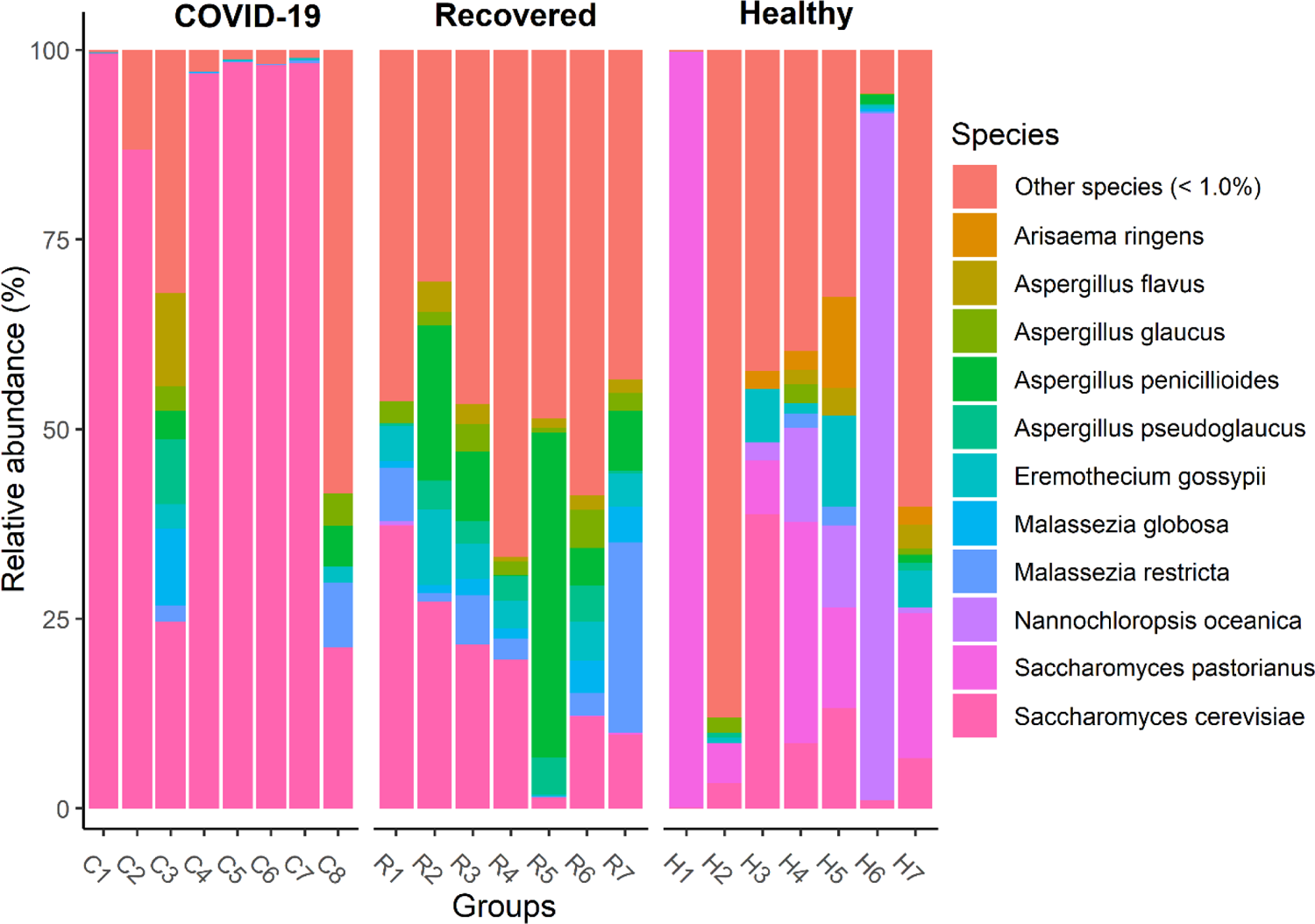
Major fungal species detected. The species-level taxonomic profile of fungal microbiomes in COVID-19 (C1-C8), Recovered (R1-R7) and Healthy (H1-H7) nasopharyngeal samples. Species with > 1% mean relative abundance are represented by different color codes against respective sample groups. Other species (< 1%) indicate the rare less abundant taxa in each group, with a mean relative abundance of < 1%. Each stacked bar plot represents the abundance of fungal species in each sample of the corresponding category. Notable differences in fungal populations are those where the species is abundant in COVID-19 and Recovered samples, and effectively not detected in the Healthy controls. The distribution and relative abundance of the fungal genera in the study metagenomes are also available in S1 Data.

Our primary microbiome compositional analysis identified 11 species as differentially abundant across Healthy, COVID-19 and Recovered metagenomes. The pairwise statistical relationship analysis of the relative abundances of these 11 taxa and health biomarkers (Healthy, SARS-CoV-2 infection and SARS-CoV-2 recovery) of the study participants showed significant variations (p ≤ 0.05, Wilcoxon test) in 11 differentially abundant fungal species (Fig 8). For instance, *N. oceanica* (p= 0.01, Wilcoxon test) and *S. pastorianus* (p= 0.05, Wilcoxon test) were the two significant and differentially abundant fungal species in Healthy controls compared to either COVID-19 patients or Recovered humans (Fig 8). The COVID-19 samples however harboured only one significantly abundant fungal species, *S. cerevisiae* (p= 0.01, Wilcoxon test) over the Healthy controls or Recovered humans. The Recovered humans however had significantly higher relative abundances of *A. penicillioides* (p= 0.01), *A. pseudoglaucus* (p= 0.01), *A. flavus* (p= 0.01), *A. fumigatus* (p= 0.05) and *E. gossypii* (p= 0.05) compared to either COVID-19 patients or Healthy controls (Fig 8). Similarly, for those species that were less abundant in three metagenomes, we also observed a stronger correlation among the microbiomes of COVID-19 patients and Recovered humans (Fig 8, S1 Data).

**Fig 8.**
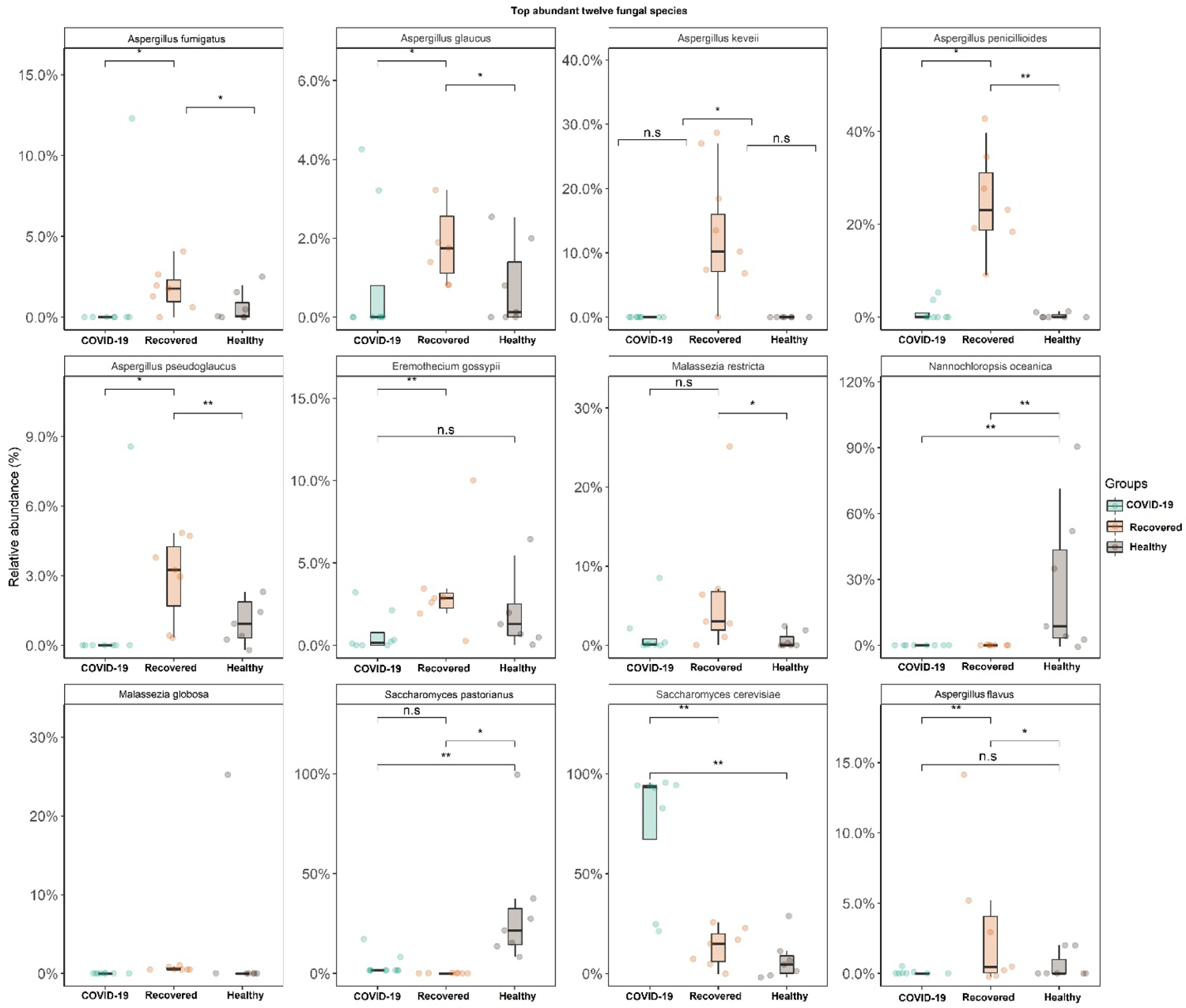
Top twelve fungal species detected. The species-level taxonomic abundance of fungal microbiomes in COVID-19, Recovered and Healthy nasopharyngeal samples. The diversity for each species is plotted on boxplots and comparisons are made with pairwise Wilcoxon test rank sum tests. Significance level (p-value) 0.0001, 0.001, 0.01, 0.05, and 0.1 are represented by the symbols “****”, “***”, “**”, “*”, and “n.s”, respectively.

### Phylogenetic relatedness of the nasopharyngeal tract fungal microbiota

The unrooted maximum likelihood tree exhibited a species-level (n=50 top abundant species) topology that was completely congruent with SARS-CoV-2 infection mediated microbiome dysbiosis (Fig 9). The resulting species tree provided high resolution of the basal relationships among microbial clades, and enabled us to evaluate the established taxonomic hierarchies. Phylogenetic analysis showed that out of 50 species, 13 descended from *Eurotiales* (26.0%) followed by 10 from *Saccharomycetales* (20.0%), four from *Auriculariales* (8.0%), one from *Eustigmatales* (2.0%) and rest of the species fall into other orders (44.0%) (Fig 9, S2 Table). Despite stronger associations among these 50 species used for phylogenetic tree reconstruction, SARS-CoV-2 infection facilitated the opportunistic inclusion of 36.54% fungal species including *S. paradoxus, S. kudriavzevii, K. lactis, K. aquatica, Y. lipolytica, C. gloeosporioides, P. album, C. sphaerospermum, F. flavus, P. hubeiensis, P. rhodozyma, Tarsonemidae sp. AD1063, C. remanei, T. govanianum,* and *G. rogosa* (Fig 9, S2 Table). Among these species, *P. rhodozyma* was found as the top abundant (10.30%) opportunistic pathogen in COVID-19 samples (S1 Data). None of these species was detected in the Healthy control samples. Simultaneously, SARS-CoV-2 inflammation was also associated with depletion of 16.0% commensal fungal population such as *E. tenuicula, P. caudatus, A. ringens, N. nucifera, A. castellanii, M. brevicollis, V. Vermiformis, A.polytricha* etc. (Fig 9, S2 Table). These species were solely present in Healthy humans nasal cavity, and not merely detected in COVID-19 patients samples Moreover, changes in microbiome composition were supported by high bootstrap values (98%–100% for each species) (Fig 9).

**Fig 9.**
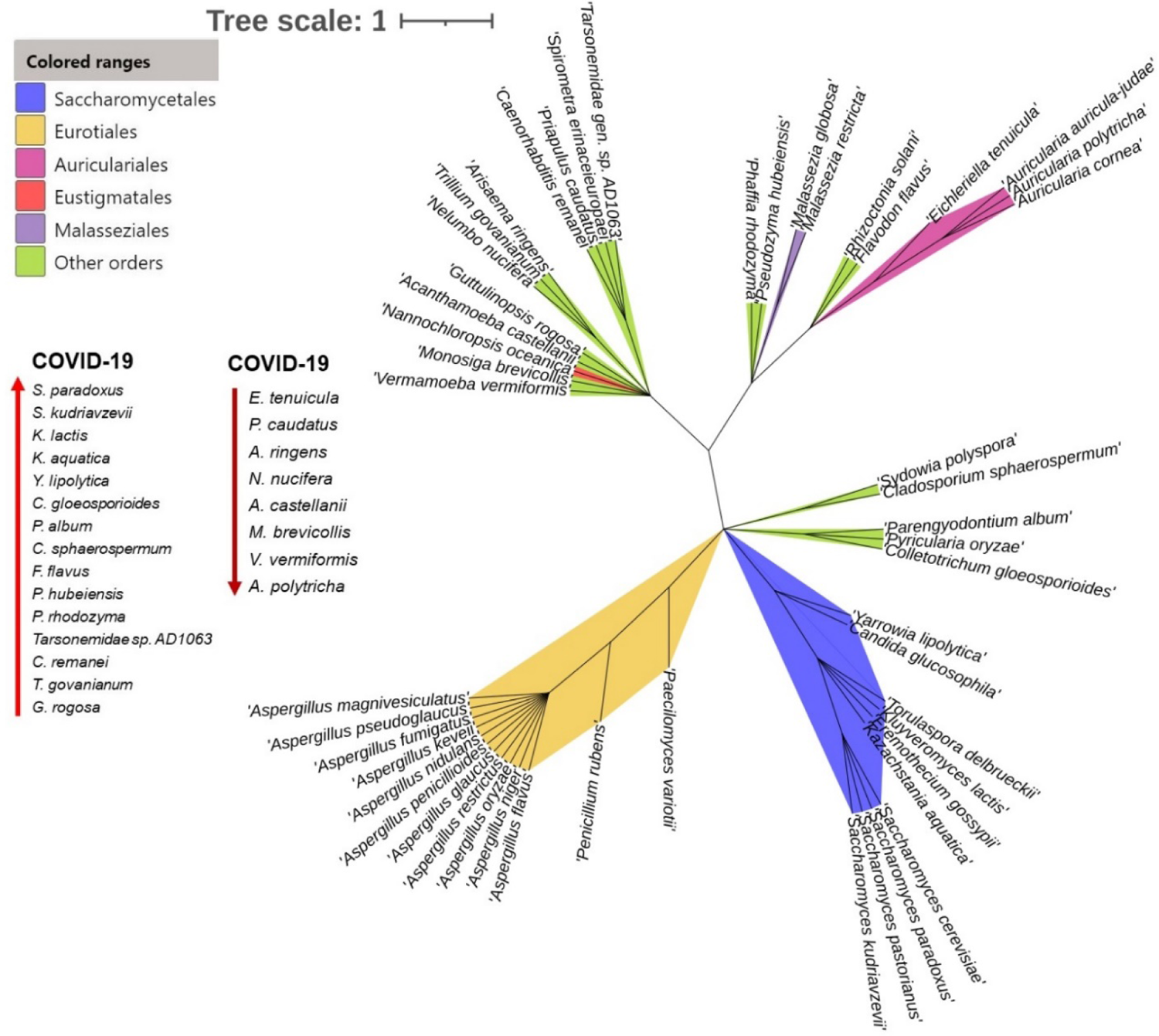
Phylogenetic relationships among detected fungal microbiomes. Unrooted phylogenetic tree showing the relationship of 50 top abundant fungal species identified in Healthy people, COVID-19 patients and Recovered humans nasopharyngeal samples. Taxonomic groups indicated by the different color ranges show the phylogenetic origin and associations of different species in the respective fungal orders, with green for *Saccharomycetales*, yellow for *Eurotiales*, pink for *Auriculariales*, red for *Eustigmatales*, purple for *Malasseziales*, and green for other orders fungal orders. The upward arrow indicates the species those had opportunistic inclusion in COVID-19 patients and not merely detected in Healthy control samples. Conversely, the downward arrow denotes the Healthy people commensal species those are dysbiosed after SARS-CoV-2 infection (absent in COVID-19 patients). Bootstrap values are calculated based on 1000 replications and the tree scale in number of substitutions per site. The fungal species in the phylogenetic tree are also available in S1 Data.

### Differentially abundant fungi show positive correlation with metabolic functional perturbations

To shed light on whether SARS-CoV-2 infection could influence the metabolic functional potentials of the concurrent fungal microbiota, we analysed the RNAseq data through KEGG pathway and SEED subsystems. By examining the correlations between the different gene families (n= 33) of the same KEGG pathway for COVID-19 patients, Recovered humans, and Healthy controls nasal microbiomes, we found significant differences (p = 0.01, Kruskal-Wallis test) in their composition and relative abundances (S1 Fig, S2 Data). Differentially abundant fungal species had significant correlations (i.e., positive or negative) with different KEGG pathways including cytokine-cytokine receptor functions, cellular processes, methane oxidation, malate dehydrogenase (mdh), oxidative phosphorylation, sulfur, protein and carbohydrates metabolism and methionine degradation have positive correlation with the dominant fungal species (Fig 10A). For instance, cytokine-cytokine receptor functions revealed strongest positive correlation with *S. cerevisiae* (Spearman correlation; r > 0.65, p < 0.001); the top abundant fungal species in COVID-19 metagenome. Likewise, cellular processes showed significant positive associations with *A. pseudoglaucus* (Spearman correlation; r > 0.6, p < 0.01) and *M. globosa* (Spearman correlation; r > 0.5, p < 0.05); two dominating fungal species in Recovered samples. In addition, *M. globosa* displayed significant positive correlations (Spearman correlation; r > 0.4, p ≤ 0.05) with methane oxidation, malate dehydrogenase (mdh), oxidative phosphorylation and sulfur metabolism (Fig 10A). Conversely, two predominating fungal species in Healthy controls, *N. oceanica* (Spearman correlation; r > 0.5, p < 0.01) and *S. pastorianus* (Spearman correlation; r > 0.5, p < 0.01) had significant negative correlation with cytokine-cytokine receptor functions, sulfur metabolism, and succinyl-CoA synthetase subunits (sucC, sucD). Similarly, *A. oryzae*, one of the prevalent fungal species in Recovered humans had strong negative correlations (Spearman correlation; r ≥ 0.5, p ≤ 0.05) with cytokine-cytokine receptor functions, sulfur metabolism and phosphotransferase system. In addition, pyruvate carboxylase (pyc), phagosome activity and focal adhesion related metabolic activities also had significant negative correlations (Spearman correlation; r ≥ 0.4, p ≤ 0.05) with the top abundant fungal species of Recovered humans (Fig 10A).

**Fig 10.**
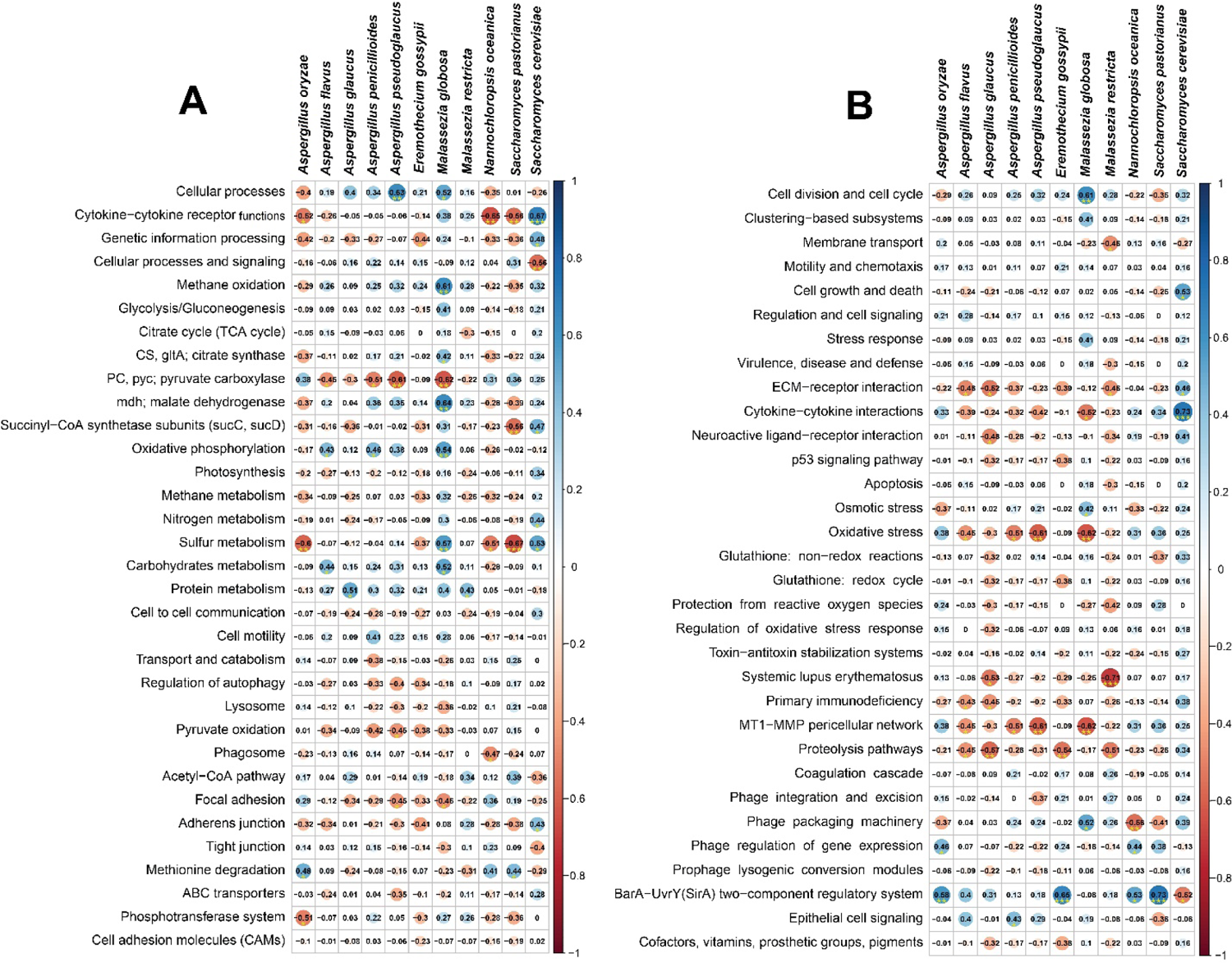
Spearman’s rank correlation matrix of the dominant fungal species and related metabolic functions. (A) Correlation between KEGG orthologues (KOs) and predominant fungal species, and (B) correlation between SEED subsystems and top abundant fungal species. The numbers display pairwise Spearman’s correlation coefficient (r). Blue and red colors indicate positive and negative correlation, respectively. The color density, circle size, and numbers reflect the scale of correlation. *Significant level (*p < 0.05; **p < 0.01; ***p < 0.001). The R packages, Hmisc (https://cran.r-project.org/web/packages/Hmisc/index.html) and corrplot (https://cran.r-project.org/web/packages/corrplot/vignettes/corrplot-intro.html) were used respectively to analyze and visualize the data.

We also sought to gain further insight into the SEED hierarchical protein functions, and found 37 statistically different (p = 0.013, Kruskal-Wallis test) subsystems in COVID (COVID-19 and Recovered) and Healthy control metagenomes. These SEED functions had significant correlations (either positive or negative) with the dominating fungal species in the respective metagenomes. As for example, the predominantly abundant fungal species in COVID-19 patients nasal swabs (*S. cerevisiae*) showed significant positive correlations with cytokine-cytokine interactions (Spearman correlation; r > 0.7, p <0.001), cell growth and death (Spearman correlation; r > 0.5, p <0.05), ECM-receptor interaction (Spearman correlation; r > 0.4, p <0.05). However, this species had also substantial negative association with BarA-UvrY(SirA) two-component regulatory system (Spearman correlation; r > 0.5, p <0.05) (Fig 10B). In contrast, BarA-UvrY(SirA) two-component regulatory system was found as the strongly correlated metabolic function in *S. pastorianus* (Spearman correlation; r > 0.7, p < 0.001), *E*. *gossypii* (Spearman correlation; r > 0.65, p < 0.001), *A. oryzae* (Spearman correlation; r > 0.55, p < 0.01) and *N. oceanica* (Spearman correlation; r > 0.5, p < 0.05). Likewise, *M. globosa* revealed significant positive correlation (Spearman correlation; r > 0.6, p < 0.01) with cell division and cell cycle related SEED functions (Fig 10B). Conversely, SEED functions including systemic lupus erythematosus, MT1-MMP pericellular network, proteolysis pathways, oxidative stress, ECM-receptor interaction and membrane transport had significant negative correlations (Spearman correlation; r ≥ 0.5, p < 0.05) with most of the top abundant fungal species in all three metagenomes (Fig 10B).

## Discussion

Emerging evidence indicated that SARS-CoV-2-infected individuals had an increased risk for coinfections. Therefore, the physicians need to be cognizant about excluding other treatable respiratory pathogens [7,10,30]. There are a great number of diverse beneficial commensal microorganisms constitutively colonizing the mucosal lining of the upper airway especially the nasopharyngeal tract (NT). These microbes comprise viruses, phages, bacteria, and fungi [7,10,31] that have elegant mutualistic relationships with the human host. In our previous study, we reported that SARS-CoV-2 infection induced dysbiosis of NT commensal bacteria, archaea and viruses with high inclusion of opportunistic pathobionts that elicite metabolic functional potential perturbations in COVID-19 patients [7]. In this study, we postulated that SARS-CoV-2 infection may also alter the NT commensal fungal population and diversity along with perturbations in their metabolic functions. To validate this hypothetical interplay between SARS-CoV-2 and resident commensal fungal microbiota in the nasal cavity of humans, we compared 22 high-throughput RNASeq data obtained from Healthy individuals, COVID-19 patients and Recovered humans.

The rapid development of automated, high-throughput sequencing methods including metagenomics, RNASeq, and bioinformatics [32] have made it possible to study the global biodiversity of fungi in various epidemiological niches including in COVI1D-9 patients. In the present study, we found a remarkable shift in the diversity and composition of the NT fungal microbiomes in COVID-19 patients and Recovered humans compared to the Healthy controls. Although, several questions remain about defining the nature of dysbiosis for any particular fungal species, our present findings showed that SARS-CoV-2 infection reduces commensal fungal population with inclusion of pathobionts in the NT of human (Fig 3 and 9). Moreover, we detected a number of microbial genomic features, altered metabolic pathways, and functional genes associated with SARS-CoV-2 pathogenesis. Although it is well established that gut microbiota has a critical role in pulmonary immunity and host’s defense against SARS-CoV-2 infection [33–35], this study for the first time determined the interactions and consequences of SARS-CoV-2 infection with resident fungal microbiome in human nasal cavity.

One of the most striking findings of the current study was the significant differences in microbiome diversity between COVID-19 patients and Healthy controls nasal cavity keeping the closest relationship of fungal population in COVID-19 patients with Recovered humans. The COVID-19 patients NT fungal microbiomes exhibited a statistically significant higher diversity (both within and between sample diversities) than those of Recovered humans and Healthy controls (Fig 1 and 2), which supports our hypothesis of dysbiosis and also agree with several recent reports [8, 36]. In contrast, most of the previous studies reported that SARS-CoV-2 infection reduces the bacterial diversity in COVID-19 patients respiratory tract and gut compared to their healthy counterpart [7, 37]. However, SARS-CoV-2 infection did not significantly alter microbiome diversity in relation to the gender of the study population.

Results from the current analysis showed that *Ascomycota* was the most predominating fungal phylum in all of three metagenomes with highest relative abundances (∼ 96.0%) in COVID-19 samples followed by Recovered humans (90.0%) and Healthy controls (88.0%). Our analysis revealed that COVID-19 patients exhibited a different composition of NT fungal microbial communities than Recovered humans and Healthy controls. Six phyla, 52 orders, 213 genera and 315 species of fungi had sole association with the SARS-CoV-2 infection (Fig 3). We found vast differences in genus-level microbiome signatures in nasal cavities of Healthy controls, COVID-19 patients and Recovered humans irrespective of the homogeneous genetic backgrounds and living status. For instance, *Nannochloropsis* had several-folds higher relative abundances in Healthy controls compared to COVID-19 and Recovered samples. Likewise, the COVID-19 patients and Recovered humans had several-folds higher abundance of *Saccharomyces* and *Aspergillus*, respectively than the Healthy controls. Moreover, the inter-individual variations in microbiome signature (fungal genera) of participants was also observed. Remarkably, more than 43.0% fungal species had sole association with Healthy humans nasal microbiomes, which were not detected in COVID-19 patients and Recovered humans indicating the potential dysbiosis of these commensal species during the pathogenic magnitudes of SARS-CoV-2 infection. These results are corroborated with our previous study where we reported that 79% commensal bacterial species found in Healthy controls were not detected in COVID-19 and Recovered humans [7]. There are lines of emerging evidences that the microbial communities in different parts of the host body can be altered in relation to pathophysiological changes [38–40]. Our present findings are also consistent with several earlier studies which reported that interactions between SARS-CoV-2 and oral microbiomes [29], and SARS-CoV-2 and gut microbiomes [41] is associated with the pathophysiology of lung diseases.

Despite hundreds of thousands of fungal species, only a few causes disease in humans. In this study, *N. oceanica* and *S. pastorianus* were two predominantly abundant fungal species in Healthy human nasopharyngeal swabs. Different species of *Nannochloropsis* are eco-sustainable bioactive microalgae which can provide human with nutritional elements including polyunsaturated fatty acids (PUFAs), polyphenols, carotenoids and vitamins [42]. Recent evidences suggested that *N. oceanica* is a chief source of eicosapentaenoic acid (EPA, 20:5n-3) and docosahexaenoic acid (DHA, 22:6 n-3) recommended for humans use due to their beneficial effects including anti-atherogenic, anti-thrombotic and anti-inflammatory properties [42, 43]. On the other hand, *S. pastorianus* is a recently evolved interspecies hybrid of *Saccharomyces* genus commonly used in the brewing industry. Different species of *Saccharomyces* are recognised as beneficial probiotics that can help humans with IBS, Crohn’s disease, diarrhoea, and a range of gastrointestinal infections [44, 45]. The co-evolution of humans and fungi suggests that complex mechanisms exist to allow the host immune system to respond to fungi [46].

One of the hallmark findings of the present study was the predominant association of *S. cerevisiae*, *P. rhodozyma* and *P. variotii* with SARS-CoV-2 infections in COVID-19 patients. A series of recent study suggested that *S. cerevisiae* should be considered as a potential opportunistic pathogen especially for patients with immunosuppression, cancer and other critical illnesses [44]. One of the latest studies demonstrated that the abundance *S. cerevisiae* in the guts of COVID-19 patients with fever were significantly higher than in COVID-19 patients with non-fever supporting our present results [47]. Bloodstream infection by *S. cerevisiae* in critically ill COVID-19 patients has recently been reported in several studies [45, 48]. *P. variotii* is considered the most prevalent agents of human infection, can affect various organ systems, primarily in immunocompromised patients or those with indwelling material [49]. Importantly, different species of *Aspergillus* were predominantly abundant in Recovered humans nasal cavity, and this shift in the fungal community is believed to be associated with re-establishment of eubiosis or a balanced NT microbiome after clearance of SARS-CoV-2. Although, *A. penicillioides* was detected as one of the top abundant fungal species in Recovered human nasal swab, this species has yet rarely been reported as a human pathogen except for cystic fibrosis in infants [50]. One of pioneering researches reported that *A. fumigatus* was the most common species causing coinfection in COVID-19 patients, followed by *A. flavus* and suggested that clinicians should keep alerting the possible occurrence of pulmonary aspergillosis in severe/critical COVID-19 patients [51, 52]. Saprophytic fungal species like *A. keveii* of the *Aspergillus* genus are the common contaminant of food and soil, their spores are ubiquitous and responsible for developing invasive aspergillosis in millions of humans each year [53]. *A. oryzae* is a low pathogenic fungus but may, like many other harmless microorganisms, grow in human tissue under exceptional circumstances [54]. In addition to *Aspergillus* spp., *S. cerevisiae* and *M. restricta* were also found to be dominating in the Recovered humans NT microbiomes. *Malassezia* spp. are lipid-dependent yeasts, inhabiting the skin and mucosa of humans and animals [55]. In adults, *M. restricta* and *M. globosa* are the major component of the healthy human skin mycobiome especially in Asia [56], and have not been linked to in the pathogenesis of any infectious diseases [57]. The interactions between human host and these fungal species can be mediated directly by specific pattern recognition receptors found on host cells and pathogen-associated molecular patterns present on fungal cell walls [56]. However, detailed clinical context is available for a very limited number of these species and data concerning their role in the pathophysiology of COVID-19 are even more scarce.

Remarkably, COVID-19 patients largely had inclusion of > 36.0% opportunistic fungal species, a part of commensal microbiota that may become pathogenic in the event of host perturbation, such as dysbiosis or immunocompromised host. *P. rhodozyma*, one of the top abundant opportunistic fungal species found in COVID-19 patients nasal cavity, is an important microorganism for its use in both the pharmaceutical industries and food industry [58]. Although, numerous studies have addressed the molecular regulatory mechanisms of cell growth and astaxanthin synthesis by *P. rhodozyma* [59, 60], however, association of this species in disease causation has not been reported yet. Different species of *Saccharomycetales* such as *S. paradoxus, S. kudriavzevii, K. lactis, K. aquatica, Y. lipolytica* had an opportunistic inclusion in COVID-19 patients (Fig 9, S1 Data). Earlier evidences suggested that different species *Saccharomycetales* can only perform opportunistic or passive crossings when epithelial barrier integrity of the NT is previously compromised by other infectious agents [70]. For instance, *S. kudriavzevii* an opportunistic pathogen especially potential hazard to the health of immunocompromised workers in the wine industry, and potentially also to consumers [61]. *S. paradoxus*, mainly found in the wild environment, is the closest relative of the domesticated yeast *S. cerevisiae*. At least five different killer toxins are produced by *S. paradoxus* which can inhibit the growth of other competing fungal species in immunocompromised host [62], and likely to cause opportunistic infection.

Despite the striking discrepancy in the phylogenetic composition and relative abundances of fungal species in three metagenomes, we found significant associations between differentially abundant fungal species and different metabolic functional pathways. Our findings revealed that enrichment of certain metabolic activities related to cytokine-cytokine receptor functions, cellular processes, methane oxidation, malate dehydrogenase, oxidative phosphorylation, sulfur, protein and carbohydrates metabolism, cell growth and death, and methionine degradation had strong positive correlation with 11 dominant fungal species, irrespective of the sample categories. The predominant fungal species of COVID-19 patients nasal cavity, *S. cerevisiae* was positively associated with cytokine-cytokine interactions, cell growth and death, and ECM-receptor interaction. These metabolic functional changes in COVID-19-associated fungal microbiomes corroborated with previously reported other respiratory viral diseases [7,63,64]. Thus, our results provided evidence that enrichment of these metabolic activities are linked to consistent shifts in the structure and composition of the NT fungal microbiome with the progression of SARS-CoV-2 pathogenesis. Similar association was also found in the dominating fungi (e.g., *A. pseudoglaucus* and *M. globose*) of Recovered humans nasal cavity, in which metabolic functions like cellular processes, methane oxidation, malate dehydrogenase, oxidative phosphorylation and sulfur metabolism were found to be positively correlated. Based on this correlative evidence, it is tempting to speculate that SARS-CoV-2 infection associated shifts in fungal microbiome in the nasal cavity might also alter the metabolic functional potentials of the related microbiomes. Spearman’s correlation analyses also revealed that the pyruvate carboxylase, phagosome activity, focal adhesion, MT1-MMP pericellular network, proteolysis pathways, oxidative stress, and membrane transport have significant negative correlations with the dominating fungal species. The metabolic health of an individual is represented by the proper functioning of organismal metabolic processes coordinated by multiple physiological systems [64]. The differentially abundant functions and pathways identified in this study corroborated with the findings from previous reports [65], and to COVID-19. However, some of the predicted metabolic features differed between COVID-19 and Healthy controls, perhaps representing metabolic changes associated with the progression of SARS-CoV-2 pathogenesis, and typical host-microbiome interactions in SARS-CoV-2 infected patients.

## Conclusion

Human nasopharyngeal microbiota plays a crucial role in providing protective responses against pathogens, particularly by regulating immune system homeostasis. Any alteration in the NT microbiota or their metabolites can cause immune dysregulation, which can impair the antiviral activity of the immune system against respiratory viruses like SARS-CoV-2. Overall, the results of this study suggested that SARS-CoV-2 infection significantly alters the diversity and population of human nasopharyngeal resident fungal microbiomes and allows the abundance of the opportunities pathogenic fungus. The distinguishable fluctuations in the fungal population and associated genomic features detected in this study might be associated with the development, treatment, and resolution of COVID-19. The SARS-CoV-2 infection results in remarkable depletion of nasopharyngeal commensal fungal population with inclusion of different opportunistic pathogens in COVID-19 and Recovered samples. Several predicted functional pathways differed between COVID-19 patient and Recovered people nasopharyngeal samples compared to Healthy individuals which reflect the roles microbial metabolic changes with the progression of SARS-CoV-2 pathogenesis. These interactions were further complicated by the common co-existence of dominant fungal microbiota that interact with both host and SARS-CoV-2. Therefore, findings of this study may serve as a benchmark for microbiome-based diagnostic markers and therapeutics for SARS-CoV-2 infection. Though, these findings support the taxonomic dysbiosis of microbiomes in the nasal cavity of COVID-19 patients, however further comprehensive study is needed to elucidate the modulation of microbiome shifts, their functional potentials and genomic expression using a larger dataset. It would be therefore imperative to carryout future research on the associations among host factors, respiratory tract commensal microbiota, and SARS-CoV-2 which will help in the development of specific therapeutic regimes against this pandemic disease.

## Materials and Methods

### Ethics statement

The protocol for sample collection from COVID-19, Recovered and Healthy humans, sample processing, transport, and RNA extraction was approved by the National Institute of Laboratory Medicine and Referral Center of Bangladesh. The study participants received a written informed consent letter consistent with the experiment.

### Subject recruitment and sample collection

We collected twenty-two (n=22) nasopharyngeal samples (including COVID-19 = 8, Recovered = 7, and Healthy = 7) from Dhaka city of Bangladesh during May to July, 2020. The suspected patients were diagnosed positive for SARS-CoV-2 infections (COVID-19) through RT-qPCR on an average 4.7 days (range: 2-9 days) after the onset of clinical signs. The confirmed patients were admitted into the dedicated COVID-19 isolation wards. These patients were tested negative for COVID-19 after 17.5 (ranged from11 to 32) days of first confirmation of SARS-CoV-2 infection, and categorized as Recovered humans (S1 Table). In this study, 68.0% and 32.0% of the selected people were male and female, respectively, and their mean age was 41.86 (ranged from 22 to 72) years. The nasopharyngeal swab samples from COVID-19 and Recovered subjects were collected on the test day (COVID-19 positive and COVID-19 negative confirmation day. The Healthy control subjects were randomly selected and these people did not show any signs and symptoms of respiratory illness. Nasopharyngeal swabs from these Healthy people were collected following the same protocol for COVID-19 and Recovered humans. The collected samples were placed in sample collection vial containing normal saline. Collected samples were preserved at − 20 °C until further use for RNA extraction and RT-qPCR assay. The RT-qPCR was performed for *ORF1ab* and *N* genes of SARS-CoV-2 using novel Coronavirus (2019-nCoV) Nucleic Acid Diagnostic Kit (PCR-Fluorescence Probing, Sansure Biotech Inc.) following the manufacturer’s instructions. The collected samples were immediately sent for RNA extraction, library preparation and sequencing [7].

### RNA extraction and sequencing

Total RNA was extracted using a PureLink viral RNA/DNA minikit (Thermo Fisher Scientific, USA). RNA was extracted from a 20 µL swab sample through lysis with sample release reagent provided by the kit and then directly used for RT-qPCR. A thermal cycling of 50 °C for 30 min was performed for reverse transcription, followed by 95 °C for 1 min, and then 45 cycles of 95 °C for 15 s, 60 °C for 30 s on an Analytik-Jena qTOWER instrument (Analytik Jena, Germany). RNAseq libraries were prepared from total RNA using the Nextera DNA Flex library preparation kit (Illumina, Inc., San Diego, CA) according to the manufacturer’s instructions where the first-strand cDNA was synthesized using SuperScript II Reverse Transcriptase (Thermo Fisher) and random primers. Paired-end (2 × 150 bp reads) sequencing of the prepared RNA library pools was performed using a NextSeq high throughput kit under Illumina platform with an Illumina NextSeq 550 sequencer at the Genomic Research Laboratory, Bangladesh Council of Scientific and Industrial Research (BCSIR), Dhaka, Bangladesh [7].

### Data processing and taxonomic identification of fungal microbiomes

The raw sequencing reads generated from Illumina platform were adapter and quality trimmed through BBDuk (with options k = 21, mink = 6, ktrim = r, ftm = 5, qtrim = rl, trimq = 20, minlen = 30, overwrite = true) [39]. Any sequence below these thresholds or reads containing more than one ‘N’ were discarded [39]. The good quality reads from COVID-19, Recovered and Healthy samples (n = 22) were analyzed using two different bioinformatics tools: the IDSeq (an open-source cloud-based pipeline to assign taxonomy) [66] and the MG-RAST (release version 4.1) (MR) and both use mapping and assembly-based hybrid method [67]. IDseq-an open-source cloud-based pipeline has been used to assign taxonomy with NT L (nucleotide alignment length in bp) >= 50 and NT %id >= 97 [7]. This pipeline used quality control, host filtering, assembly-based alignment and taxonomic reporting aligning to NCBI nucleotide database. In MR analysis, the uploaded reads were subjected to optional quality filtering with dereplication and host DNA removal, and finally functional assignment. For this pipeline, we employed the ‘‘Best Hit Classification’’ option to determine taxonomic abundance using the NCBI database as a reference with the following set parameters: maximum *e*-value of 1×10^-30^; minimum identity of 90% using a minimum alignment length of 20 as the set parameters. Microbial taxa that were detected in one group of sample but not detected in rest of the two groups are denoted as solely (unique) associated microbiomes [39]. We simultaneously uploaded the filtered sequence data in both pipelines with proper embedded metadata.

### Functional profiling of the microbiomes

We performed the fungal metabolic functional classification through mapping the reads onto the Kyoto Encyclopaedia of Genes and Genomes (KEGG) database [68], and SEED subsystem identifiers [67] of the MR server using the partially modified set parameters (*e-*value cut off: 1×10^-30^, min. % identity cut off: 90%, and min. alignment length cut off: 20) [69]. The association between metabolic functions and dominant fungal species were measured using the Spearman’s correlation coefficient and significance tests [70]. The R packages, Hmisc (https://cran.r-project.org/web/packages/Hmisc/index.html) and corrplot (https://cran.r-project.org/web/packages/corrplot/vignettes/corrplot-intro.html) were used respectively to analyze and visualize the data.

### Statistical analysis

Read normalization in each sample was performed using median sequencing depth through Phyloseq (version 4.1) package in R [71]. We calculated the alpha diversity (diversity within samples) using the observed species, Chao1, ACE, Shannon, Simpson and InvSimpson diversity indices, and performed the non-parametric Wilcoxon rank-sum test to evaluate diversity differences in different samples. A principal coordinate analysis (PCoA) based on the Bray-Curtis distance method was performed to visualize differences in fungal diversity across three metagenomes. To calculate the significance of variability patterns of the microbiomes (generated between sample categories), we performed PERMANOVA (permutational multivariate analysis of variance) using 999 permutations on all three sample types at the same time and compared them pairwise. For these statistical analyses, pairwise non-parametric Wilcoxon test was performed using the Phyloseq and Vegan (package 2.5.1 of R 3.4.2) programs [72]. Dominant fungal community were determined with ≥ 1% median relative abundance by groups. After filtering, 11 taxa remained for which Spearman’s correlation analysis between KEGG pathways and SEED functions pathways was done in Hmisc’s rcorr function [73] and the corrplot function [74] of the corrplot R package as mentioned in the previous section. In addition, Kruskal-Wallis test was also applied at different KEGG and SEED subsystems levels through IBM SPSS (SPSS, Version 23.0, IBM Corp., NY, USA) [39].

## Supplementary Figure with Legend

**S1 Fig.**
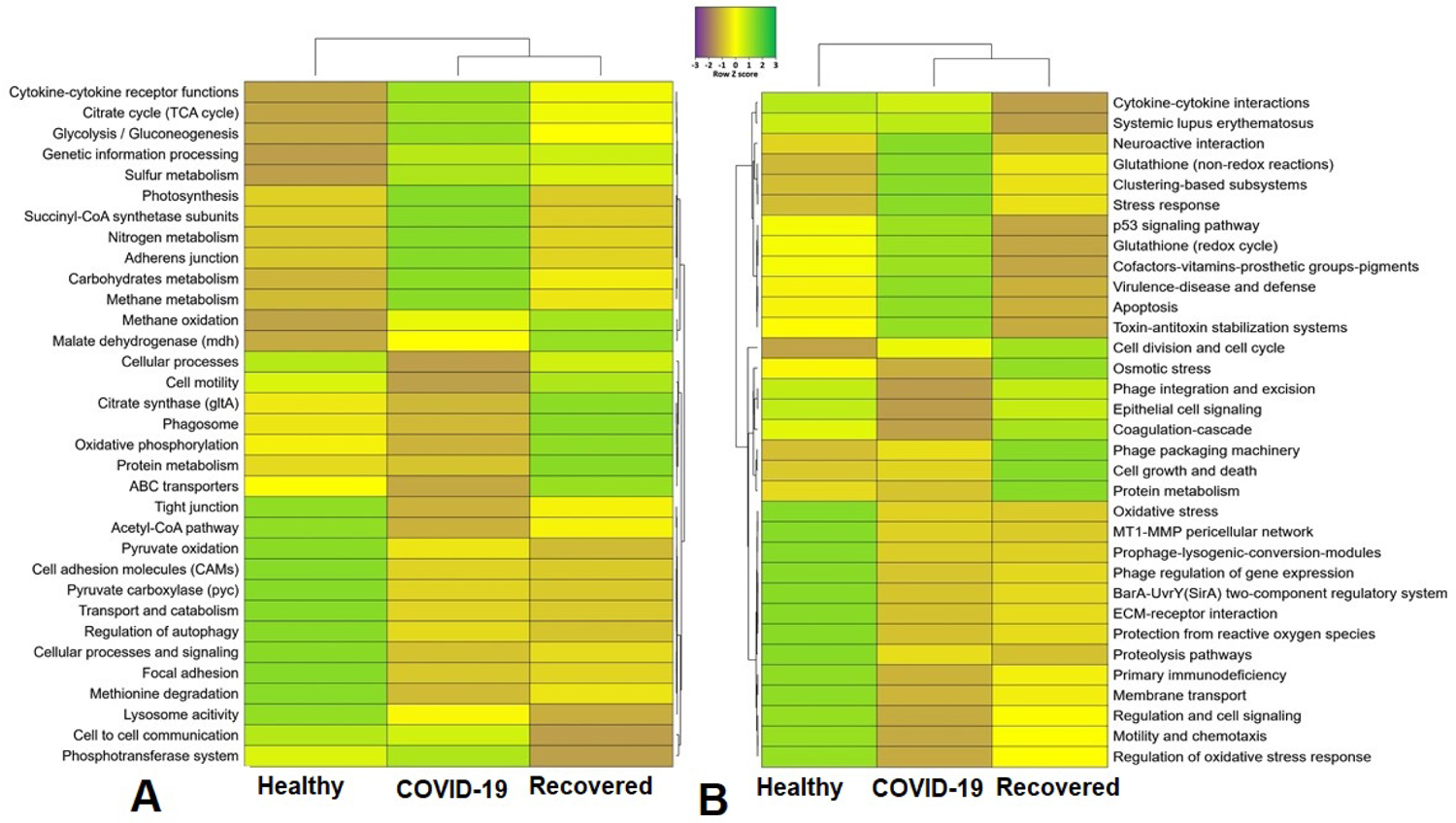
Metabolic functional potentials of the fungal microbiota. Heatmap showing (A) KEGG orthologues (KOs) and (B) SEED subsystems associated with fungal metabolism in Healthy, COVID-19 and Recovered metagenomes.

## Funding

This study received no financial support.

## Competing interests

The authors declare no competing interests.

## Data availability

The sequence data reported in this article has been deposited in the National Center for Biotechnology Information (NCBI) under BioProject accession number PRJNA720904.

## Acknowledgements

The authors would like to thank those who provided us the samples.

## Supplementary information

Supplementary information supporting the results of the study are available in this article as S1 Fig, S1-S2 Table and S1-S2 Data.

## Notes

### Competing Interest Statement

The authors have declared no competing interest.

## References

1. Grau-Expósito J, Perea D, Suppi M, Massana N, Vergara A, et al. (2022) Evaluation of SARS-CoV-2 entry, inflammation and new therapeutics in human lung tissue cells. PLoS pathogens 18: e1010171.

2. Hoque MN, Chaudhury A, Akanda MAM, Hossain MA, Islam MT (2020) Genomic diversity and evolution, diagnosis, prevention, and therapeutics of the pandemic COVID-19 disease. PeerJ 8: e9689.

3. Andersen KG, Rambaut A, Lipkin WI, Holmes EC, Garry RF (2020) The proximal origin of SARS-CoV-2. Nature medicine 26: 450–452.

4. Rafiqul Islam S, Foysal MJ, Hoque MN, Mehedi H, Salauddin A, et al. Dysbiosis of oral and gut microbiomes in SARS-CoV-2 infected patients in Bangladesh: elucidating the role of opportunistic gut microbes. Frontiers in Medicine: 163.

5. Gallo O, Locatello LG, Mazzoni A, Novelli L, Annunziato F (2020) The central role of the nasal microenvironment in the transmission, modulation, and clinical progression of SARS-CoV-2 infection. Mucosal immunology: 1–12.

6. Li X, Ma X (2020) Acute respiratory failure in COVID-19: is it “typical” ARDS? Critical Care 24: 1–5.

7. Hoque MN, Sarkar MMH, Rahman MS, Akter S, Banu TA, et al. (2021) SARS-CoV-2 infection reduces human nasopharyngeal commensal microbiome with inclusion of pathobionts. Scientific Reports 11: 24042.

8. Hoque MN, Rahman MS, Ahmed R, Hossain MS, Islam MS, et al. (2021) Diversity and genomic determinants of the microbiomes associated with COVID-19 and non-COVID respiratory diseases. Gene Reports 23: 101200.

9. Trypsteen W, Van Cleemput J, Snippenberg Wv, Gerlo S, Vandekerckhove L (2020) On the whereabouts of SARS-CoV-2 in the human body: A systematic review. PLoS pathogens 16: e1009037.

10. Hoque MN, Akter S, Mishu ID, Islam MR, Rahman MS, et al. (2021) Microbial co-infections in COVID-19: Associated microbiota and underlying mechanisms of pathogenesis. Microbial Pathogenesis: 104941.

11. Feldman C, Anderson R (2021) The role of co-infections and secondary infections in patients with COVID-19. Pneumonia 13: 1–15.

12. Bardi T, Pintado V, Gomez-Rojo M, Escudero-Sanchez R, Lopez AA, et al. (2021) Nosocomial infections associated to COVID-19 in the intensive care unit: clinical characteristics and outcome. European Journal of Clinical Microbiology & Infectious Diseases 40: 495–502.

13. Zhang G, Hu C, Luo L, Fang F, Chen Y, et al. (2020) Clinical features and short-term outcomes of 221 patients with COVID-19 in Wuhan, China. Journal of Clinical Virology 127: 104364.

14. Yue H, Zhang M, Xing L, Wang K, Rao X, et al. (2020) The epidemiology and clinical characteristics of co-infection of SARS-CoV-2 and influenza viruses in patients during COVID-19 outbreak. Journal of medical virology 92: 2870–2873.

15. Cuadrado-Payán E, Montagud-Marrahi E, Torres-Elorza M, Bodro M, Blasco M, et al. (2020) SARS-CoV-2 and influenza virus co-infection. Lancet (London, England) 395: e84.

16. Antinori S, Galimberti L, Milazzo L, Ridolfo AL (2020) Bacterial and fungal infections among patients with SARS-CoV-2 pneumonia. hospitals 85: 90-96.

17. Gangneux J-P, Dannaoui E, Fekkar A, Luyt C-E, Botterel F, et al. (2021) Fungal infections in mechanically ventilated patients with COVID-19 during the first wave: the French multicentre MYCOVID study. The Lancet Respiratory Medicine.

18. Baddley JW, Thompson III GR, Chen SC-A, White PL, Johnson MD, et al. Coronavirus Disease 2019– Associated Invasive Fungal Infection; 2021. Oxford University Press US. pp. ofab510.

19. Schauwvlieghe AF, Rijnders BJ, Philips N, Verwijs R, Vanderbeke L, et al. (2018) Invasive aspergillosis in patients admitted to the intensive care unit with severe influenza: a retrospective cohort study. The Lancet Respiratory Medicine 6: 782–792.

20. Choudhary NK, Jain AK, Soni R, Gahlot N (2021) Mucormycosis: A deadly black fungus infection among COVID-19 patients in India. Clinical epidemiology and global health 12: 100900.

21. Soni S, Namdeo Pudake R, Jain U, Chauhan N (2022) A systematic review on SARS-CoV-2-associated fungal coinfections. Journal of Medical Virology 94: 99–109.

22. Aranjani JM, Manuel A, Abdul Razack HI, Mathew ST (2021) COVID-19–associated mucormycosis: Evidence-based critical review of an emerging infection burden during the pandemic’s second wave in India. PLoS neglected tropical diseases 15: e0009921.

23. Chen N, Zhou M, Dong X, Qu J, Gong F, et al. (2020) Epidemiological and clinical characteristics of 99 cases of 2019 novel coronavirus pneumonia in Wuhan, China: a descriptive study. The lancet 395: 507-513.

24. Song G, Liang G, Liu W (2020) Fungal co-infections associated with global COVID-19 pandemic: a clinical and diagnostic perspective from China. Mycopathologia: 1–8.

25. Thevissen K, Jacobs C, Holtappels M, Toda M, Verweij P, et al. (2020) International survey on influenza-associated pulmonary aspergillosis (IAPA) in intensive care units: responses suggest low awareness and potential underdiagnosis outside Europe. Critical Care 24: 1–5.

26. Lai C-C, Yu W-L (2020) COVID-19 associated with pulmonary aspergillosis: A literature review. Journal of Microbiology, Immunology and Infection.

27. Mirzaei R, Goodarzi P, Asadi M, Soltani A, Aljanabi HAA, et al. (2020) Bacterial co-infections with SARS-CoV-2. IUBMB life 72: 2097–2111.

28. Quah J, Jiang B, Tan PC, Siau C, Tan TY (2018) Impact of microbial Aetiology on mortality in severe community-acquired pneumonia. BMC infectious diseases 18: 1–9.

29. Bao L, Zhang C, Dong J, Zhao L, Li Y, et al. (2020) Oral microbiome and SARS-CoV-2: beware of lung co-infection. Frontiers in microbiology 11: 1840.

30. Musuuza JS, Watson L, Parmasad V, Putman-Buehler N, Christensen L, et al. (2021) Prevalence and outcomes of co-infection and superinfection with SARS-CoV-2 and other pathogens: a systematic review and meta-analysis. PloS one 16: e0251170.

31. Soltani S, Zakeri A, Zandi M, Kesheh MM, Tabibzadeh A, et al. (2021) The role of bacterial and fungal human respiratory microbiota in COVID-19 patients. BioMed research international 2021.

32. Huffnagle GB, Noverr MC (2013) The emerging world of the fungal microbiome. Trends in microbiology 21: 334–341.

33. Dhar D, Mohanty A (2020) Gut microbiota and Covid-19-possible link and implications. Virus research: 198018.

34. Yeoh YK, Zuo T, Lui GC-Y, Zhang F, Liu Q, et al. (2021) Gut microbiota composition reflects disease severity and dysfunctional immune responses in patients with COVID-19. Gut 70: 698–706.

35. Sencio V, Machado MG, Trottein F (2021) The lung–gut axis during viral respiratory infections: The impact of gut dysbiosis on secondary disease outcomes. Mucosal Immunology: 1–9.

36. Moreira-Rosário A, Marques C, Pinheiro H, Araújo JR, Ribeiro P, et al. (2021) Gut microbiota diversity and C-Reactive Protein are predictors of disease severity in COVID-19 patients. bioRxiv.

37. Engen PA, Naqib A, Jennings C, Green SJ, Landay A, et al. (2021) Nasopharyngeal Microbiota in SARS-CoV-2 Positive and Negative Patients. Biological procedures online 23: 1–6.

38. Scepanovic P, Hodel F, Mondot S, Partula V, Byrd A, et al. (2019) A comprehensive assessment of demographic, environmental, and host genetic associations with gut microbiome diversity in healthy individuals. Microbiome 7: 1–15.

39. Hoque MN, Istiaq A, Clement RA, Sultana M, Crandall KA, et al. (2019) Metagenomic deep sequencing reveals association of microbiome signature with functional biases in bovine mastitis. Scientific reports 9: 1–14.

40. Hoque M, Sultana M, Hossain A (2021) Dynamic Changes in Microbiome Composition and Genomic Functional Potentials in Bovine Mastitis. J Data Mining Genomics Proteomics 12: 232.

41. Kalantar-Zadeh K, Ward SA, Kalantar-Zadeh K, El-Omar EM (2020) Considering the effects of microbiome and diet on SARS-CoV-2 infection: nanotechnology roles. ACS nano 14: 5179–5182.

42. Alves SP, Mendonça SH, Silva JL, Bessa RJ (2018) Nannochloropsis oceanica, a novel natural source of rumen-protected eicosapentaenoic acid (EPA) for ruminants. Scientific reports 8: 1–10.

43. Vítor A, Francisco AE, Silva J, Pinho M, Huws SA, et al. (2021) Freeze-dried Nannochloropsis oceanica biomass protects eicosapentaenoic acid (EPA) from metabolization in the rumen of lambs. Scientific reports 11: 1–16.

44. Algazaq JN, Akrami K, Martinez F, McCutchan A, Bharti AR (2017) Saccharomyces cerevisiae laryngitis and oral lesions in a patient with laryngeal carcinoma. Case reports in infectious diseases 2017.

45. Ventoulis I, Sarmourli T, Amoiridou P, Mantzana P, Exindari M, et al. (2020) Bloodstream infection by Saccharomyces cerevisiae in two COVID-19 patients after receiving supplementation of Saccharomyces in the ICU. Journal of Fungi 6: 98.

46. Romani L (2011) Immunity to fungal infections. Nature Reviews Immunology 11: 275–288.

47. Zhou Y, Shi X, Fu W, Xiang F, He X, et al. (2021) Gut Microbiota Dysbiosis Correlates with Abnormal Immune Response in Moderate COVID-19 Patients with Fever. Journal of Inflammation Research 14: 2619.

48. Hoenigl M (2021) Invasive fungal disease complicating coronavirus disease 2019: when it rains, it spores. Oxford University Press US. pp. e1645–e1648.

49. Sprute R, Salmanton-García J, Sal E, Malaj X, Falces-Romero I, et al. (2021) Characterization and outcome of invasive infections due to Paecilomyces variotii: analysis of patients from the FungiScope® registry and literature reports. Journal of Antimicrobial Chemotherapy 76: 765–774.

50. Gupta K, Gupta P, Mathew JL, Bansal A, Singh G, et al. (2016) Fatal Disseminated Aspergillus Penicillioides Infection in a 3-Month-Old Infant with Suspected Cystic Fibrosis: Autopsy Case Report with Review of Literature. Pediatric and Developmental Pathology 19: 506–511.

51. Lai C-C, Yu W-L (2021) COVID-19 associated with pulmonary aspergillosis: A literature review. Journal of Microbiology, Immunology and Infection 54: 46–53.

52. Zuo T, Zhang F, Lui GC, Yeoh YK, Li AY, et al. (2020) Alterations in gut microbiota of patients with COVID-19 during time of hospitalization. Gastroenterology 159: 944–955. e948.

53. Gautier M, Normand A-C, Ranque S (2016) Previously unknown species of Aspergillus. Clinical Microbiology and Infection 22: 662–669.

54. Barbesgaard P, Heldt-Hansen HP, Diderichsen B (1992) On the safety of Aspergillus oryzae: a review. Applied microbiology and biotechnology 36: 569–572.

55. Rhimi W, Theelen B, Boekhout T, Otranto D, Cafarchia C (2020) Malassezia spp. yeasts of emerging concern in fungemia. Frontiers in cellular and infection microbiology 10: 370.

56. Krzyściak P, Bakuła Z, Gniadek A, Garlicki A, Tarnowski M, et al. (2020) Prevalence of Malassezia species on the skin of HIV-seropositive patients. Scientific Reports 10: 1–13.

57. Wu G, Zhao H, Li C, Rajapakse MP, Wong WC, et al. (2015) Genus-wide comparative genomics of Malassezia delineates its phylogeny, physiology, and niche adaptation on human skin. PLoS genetics 11: e1005614.

58. Elwan H, Elnesr S, Abdallah Y, Hamdy A, El-Bogdady A (2019) Red yeast (Phaffia rhodozyma) as a source of Astaxanthin and its impacts on productive performance and physiological responses of poultry. World’s Poultry Science Journal 75: 273–284.

59. Satoh A, Tsuji S, Okada Y, Murakami N, Urami M, et al. (2009) Preliminary clinical evaluation of toxicity and efficacy of a new astaxanthin-rich Haematococcus pluvialis extract. Journal of Clinical Biochemistry and Nutrition 44: 280–284.

60. Miao L, Chi S, Wu M, Liu Z, Li Y (2019) Deregulation of phytoene-β-carotene synthase results in derepression of astaxanthin synthesis at high glucose concentration in Phaffia rhodozyma astaxanthin-overproducing strain MK19. BMC microbiology 19: 1–11.

61. Douglass AP, Offei B, Braun-Galleani S, Coughlan AY, Martos AA, et al. (2018) Population genomics shows no distinction between pathogenic Candida krusei and environmental Pichia kudriavzevii: one species, four names. PLoS pathogens 14: e1007138.

62. Fredericks LR, Lee MD, Crabtree AM, Boyer JM, Kizer EA, et al. (2021) The species-specific acquisition and diversification of a K1-like family of killer toxins in budding yeasts of the Saccharomycotina. PLoS genetics 17: e1009341.

63. Wu J, Wang K, Wang X, Pang Y, Jiang C (2020) The role of the gut microbiome and its metabolites in metabolic diseases. Protein & Cell: 1–14.

64. Ayres JS (2020) A metabolic handbook for the COVID-19 pandemic. Nature metabolism 2: 572–585.

65. Breiman A, Ruvën-Clouet N, Le Pendu J (2020) Harnessing the natural anti-glycan immune response to limit the transmission of enveloped viruses such as SARS-CoV-2. PLoS Pathogens 16: e1008556.

66. Kalantar KL, Carvalho T, de Bourcy CF, Dimitrov B, Dingle G, et al. (2020) IDseq—an open source cloud-based pipeline and analysis service for metagenomic pathogen detection and monitoring. Gigascience 9: giaa111.

67. Glass EM, Wilkening J, Wilke A, Antonopoulos D, Meyer F (2010) Using the metagenomics RAST server (MG-RAST) for analyzing shotgun metagenomes. Cold Spring Harbor Protocols 2010: pdb. prot5368.

68. Kanehisa M, Sato Y, Furumichi M, Morishima K, Tanabe M (2019) New approach for understanding genome variations in KEGG. Nucleic acids research 47: D590–D595.

69. Hoque MN, Istiaq A, Clement RA, Gibson KM, Saha O, et al. (2020) Insights into the resistome of bovine clinical mastitis microbiome, a key factor in disease complication. Frontiers in Microbiology 11: 860.

70. Rahman MS, Hoque MN, Puspo JA, Islam MR, Das N, et al. (2021) Microbiome signature and diversity regulates the level of energy production under anaerobic condition. Scientific reports 11: 1–23.

71. McMurdie PJ, Holmes S (2013) phyloseq: an R package for reproducible interactive analysis and graphics of microbiome census data. PloS one 8: e61217.

72. Beck J, Holloway JD, Schwanghart W (2013) Undersampling and the measurement of beta diversity. Methods in Ecology and Evolution 4: 370–382.

73. Harrell Jr F, Harrell Jr M (2019) Package ‘hmisc’. CRAN2018 2019.

74. Wei T, Simko V, Levy M, Xie Y, Jin Y, et al. (2017) Package ‘corrplot’. Statistician 56: e24.

